# Axes of inter-sample variability among transcriptional neighborhoods reveal disease-associated cell states in single-cell data

**DOI:** 10.1101/2021.04.19.440534

**Authors:** Yakir Reshef, Laurie Rumker, Joyce B. Kang, Aparna Nathan, Ilya Korsunsky, Samira Asgari, Megan B. Murray, D. Branch Moody, Soumya Raychaudhuri

## Abstract

As single-cell datasets grow in sample size, there is a critical need to characterize cell states that vary across samples and associate with sample attributes like clinical phenotypes. Current statistical approaches typically map cells to cell-type clusters and examine sample differences through that lens alone. Here we present covarying neighborhood analysis (CNA), an unbiased method to identify cell populations of interest with greater flexibility and granularity. CNA characterizes dominant axes of variation across samples by identifying groups of very small regions in transcriptional space—termed neighborhoods—that covary in abundance across samples, suggesting shared function or regulation. CNA can then rigorously test for associations between any sample-level attribute and the abundances of these covarying neighborhood groups. We show in simulation that CNA enables more powerful and accurate identification of disease-associated cell states than a cluster-based approach. When applied to published datasets, CNA captures a Notch activation signature in rheumatoid arthritis, redefines monocyte populations expanded in sepsis, and identifies a previously undiscovered T-cell population associated with progression to active tuberculosis.

## Introduction

High-dimensional profiling of single cells is a central tool for understanding complex biological systems[1]. Cells gathered from distinct samples are used to characterize cell states that associate with a sample attribute like a clinical phenotype or experimental perturbation. Current methods for analyzing multi-sample single-cell datasets typically impose a global transcriptional structure on the dataset by partitioning cells into groups through clustering[2, 3]. The data are then analyzed solely through this lens by asking whether a sample attribute is associated with expansion or depletion of any clusters. Such approaches assume the underlying biology is well captured by the imposed structure and often require substantial tuning of parameters such as clustering resolution[4].

Here we present covarying neighborhood analysis (CNA), a method for characterizing dominant axes of inter-sample variability and conducting association testing in single-cell datasets without requiring a pre-specified transcriptional structure. The core notion of CNA is the value of granular analysis of neighborhoods—very small regions in transcriptional space—with aggregation of neighborhoods according to their covariance across samples. We posit that groups of neighborhoods that change in abundance together across samples are likely to represent biologically meaningful units that share function, regulatory influences, or both. CNA can be used to define these covarying neighborhood groups and then identify statistical associations between them and any sample-level attribute. One published method, MELD, has already demonstrated the potential of neighborhood-scale abundance information in datasets with small sample size[5]; however, this method does not provide a framework for determining statistical significance in order to differentiate true from false discoveries. As we show, the large number of neighborhoods in many single-cell datasets makes well-powered association testing at this granularity a challenge. CNA addresses this challenge by leveraging the extensive covariance structure that we show exists across neighborhoods. As a result, CNA offers both a data-dependent, parsimonious representation of single-cell data and well-powered and accurate association testing.

By testing simulated sample attributes in real single-cell data, we demonstrate that CNA is well calibrated and, compared to cluster-based analysis, detects diverse signals with improved power and accuracy. We then apply CNA to three published datasets[6-8], demonstrating that it both refines and expands upon the associations previously found using standard approaches.

## Results

### Overview of Methods

Covarying neighborhood analysis (CNA) relies on a representation of each sample in a single-cell dataset by its abundance of cells across neighborhoods. Starting with a cell-cell similarity graph, we define one neighborhood per cell *m* in the dataset: every other cell *m*′ belongs to the neighborhood anchored at cell *m* according to the probability that a random walk in the graph from *m*′ will arrive at *m* after *s* steps (**Methods**; **Figure 1A**). CNA chooses *s* in a data-dependent manner to minimize neighborhood size while ensuring that neighborhoods are not dominated by cells from only a few samples. **Supplementary Figure 1** and **Supplementary Table 1** show example neighborhoods and average neighborhood sizes for the real datasets analyzed in this paper.

**Figure 1:**
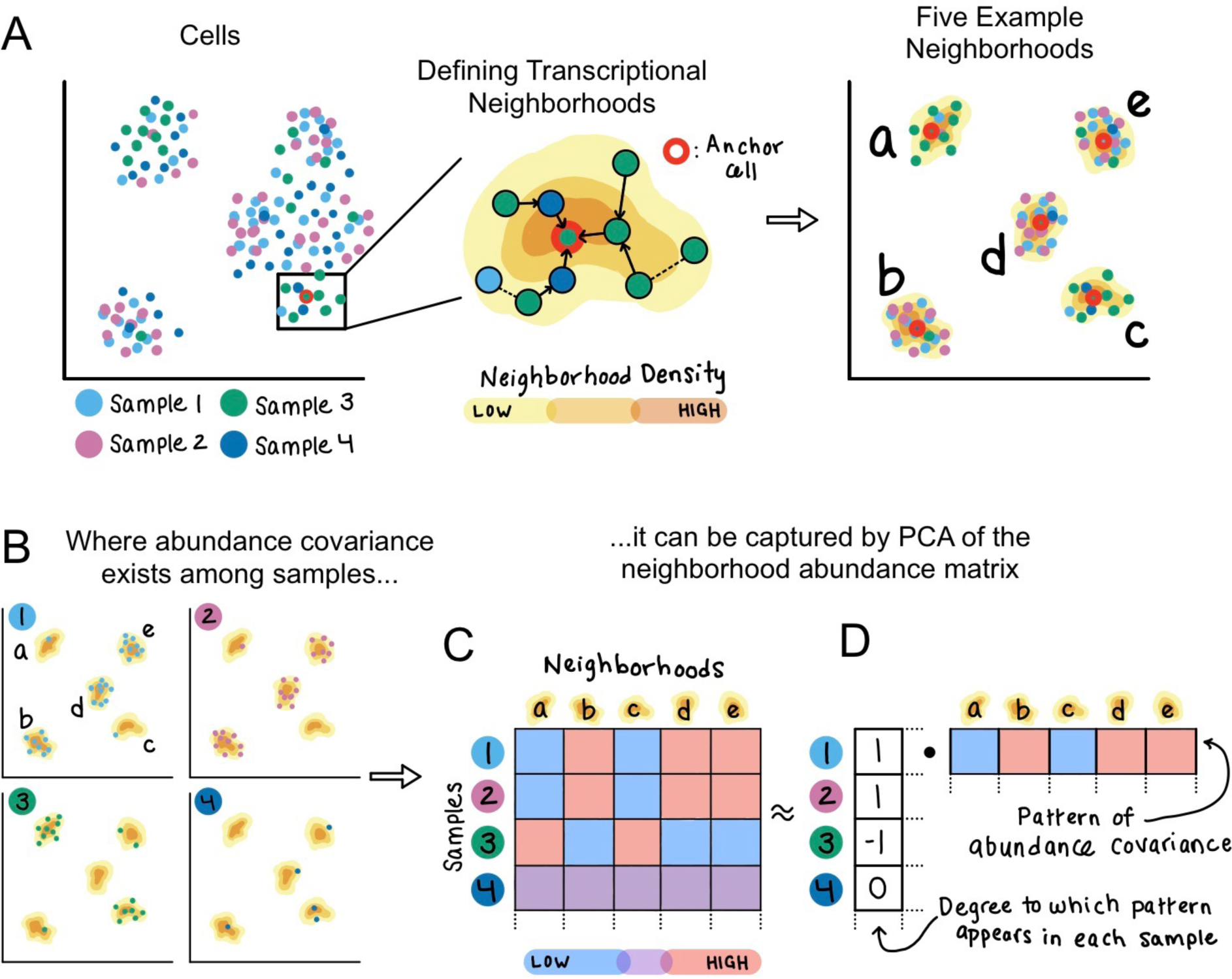
Method schematic. **(A**) Given an example dataset of single cells sampled from four individuals, CNA defines one transcriptional neighborhood per cell in the dataset. Each other cell in the dataset belongs to this neighborhood according to the probability that a random walk in the cell-cell similarity graph from that cell will arrive at the neighborhood’s anchor cell after a certain number of steps. Five example neighborhoods a-e are depicted. (**B**) Examining the representation of cells from each sample in these example neighborhoods reveals a pattern of abundance covariation. Neighborhoods b, d, and e tend to have a high abundance when neighborhoods a and e have low abundance, and vice versa. This covariation pattern appears in samples 1-3 but not in sample 4. (**C**) The neighborhood abundance matrix (NAM) quantifies the fractional abundance of cells in each neighborhood for each sample; we indicate higher abundance with red and lower abundance with blue. (**D**) Dominant patterns of abundance covariation across neighborhoods can be illuminated by factorizing the NAM, for example with PCA. The principal component corresponding to this example has per-neighborhood loadings that capture the neighborhood covariance pattern, as well as per-sample loadings that reflect the degree to which the covariance pattern appears in each sample.

We aggregate this information into a *neighborhood abundance matrix* (NAM) whose *n, m*-th entry is the relative abundance of cells from sample *n* in neighborhood *m* (**Figure 1B-C**). We then apply principal components analysis to the NAM to define neighborhood groups whose abundances change in concert across samples (**Figure 1D**). For each NAM principal component (NAM-PC), the neighborhoods with positive loadings tend to have high abundance together in the samples for which the neighborhoods with negative loadings have low abundance. Likewise, the sample loadings for each NAM-PC yield information about the extent to which that NAM-PC’s pattern of covarying neighborhoods appears in each sample.

NAM-PCs can be used to characterize transcriptional changes that comprise the axes of greatest variation in neighborhood abundances across samples. They can also be used to test for associations between these transcriptional changes and a per-sample attribute of interest, *e*.*g*., a clinical attribute, genotype, or experimental condition. To perform this test, we model the attribute value for each sample as a linear function of the sample’s loadings on the first *k* NAM-PCs, where *k* is chosen in a data-dependent manner to optimize model performance without overfitting (**Methods**). We report a p-value for this association by permuting attribute values within experimental batches to obtain a null distribution.

Finally, we define the specific cell populations driving any detected associations. We do so by using the neighborhood loadings on the first *k* NAM-PCs and the estimated per-PC effect sizes from our linear model to estimate per-neighborhood correlations between neighborhood abundance—as captured by the first *k* NAM-PCs—and the sample attribute (**Methods**). We report false discovery rates (FDRs) for each per-neighborhood association by again permuting attribute values within experimental batches to obtain null distributions. We refer to the abundance correlation between the attribute and the neighborhood anchored at each cell as the *neighborhood coefficient* of that cell. We control for sample-level confounders, such as demographic variables, technical parameters and batch effects, by linearly projecting them out of the NAM and the attribute prior to association testing (**Methods**). We have released open-source software implementing the method (**URLs**).

CNA requires no parameter tuning, and it has favorable runtime properties: given a nearest-neighbor graph, computing the NAM and conducting permutation-based association testing takes <1 minute (and 579MB memory) for a dataset of >500,000 cells and >250 samples.

### Performance assessment with simulations

We used real single-cell data and simulated per-sample attributes to assess CNA’s calibration (type I error) and to compare CNA’s statistical power (type II error) against cluster-based analysis. This published dataset of 259 patients previously infected with *Mycobacterium tuberculosis* contains 500,089 memory T cells in a canonical correlation analysis (CCA)-based per-cell joint representation of whole-transcriptome mRNA and abundances of 31 surface proteins[6]. In addition to assessing calibration and power, we also assessed CNA’s accuracy in identifying the precise cell populations underlying an association by computing the correlation between the per-cell ground-truth values used to create the simulated attribute and effect sizes estimated by the method (**Methods**; **Supplementary Figure 2**).

To assess type I error, we simulated sample attributes without true associations to the data and found CNA was well-calibrated. We first permuted patient age across all samples and observed a p<0.05 global association in 41/1000 trials (type I error rate at α=0.05 of 0.041±0.013; **Supplementary Figure 3**). We next permuted patient ages within experimental batches and observed p<0.05 for 44/1000 trials (type I error 0.044±0.013; **Supplementary Figure 3**). Finally, to simulate extreme batch effects, we selected batches at random and for each randomly selected batch assigned case status to the samples in the selected batch and control status to all other samples. We observed p<0.05 for 60/1000 trials (type I error 0.060±0.015; **Supplementary Figure 3**).

To assess CNA’s power and accuracy, we simulated sample attributes with true associations to different types of cell populations and compared CNA’s performance to that of a cluster-based association test using Mixed-effects modeling of Associations of Single Cells (MASC)[9]; MASC offers greater power than a t-test or linear model by accounting for per-cell information[10]. For CNA, power was defined as the proportion of simulations with global p<0.05. For MASC, power was defined as the proportion of simulations for which at least one cluster achieved p<0.05/[total clusters]. Cluster-based analysis is sensitive to the choice of parameters such as the resolution parameter, and users typically explore a range of resolutions before selecting one[4]. To reflect this, we ran MASC using four different clustering resolutions. We aggregated power results across these resolutions by taking the minimum p-value and correcting for the four resolutions tested. We aggregated accuracy results by taking the average accuracy across the tested resolutions (**Methods**).

We simulated three signal types, each at a variety of noise levels: 1) cluster abundance, where the attribute is a sample’s abundance of cells from a given cluster (matching the cluster-based analysis model; **Figure 2A**); 2) global gene expression program (GEP), where the attribute is a sample’s average use of a GEP across all cells (**Figure 2B**); and 3) cluster-specific GEP, where the attribute is a sample’s average use of a GEP across cells in one cluster (**Figure 2C**). We used principal components computed from the matrix of cells-by-canonical variables for the whole dataset or for cells within a cluster as our global and cluster-specific GEPs, respectively.

**Figure 2:**
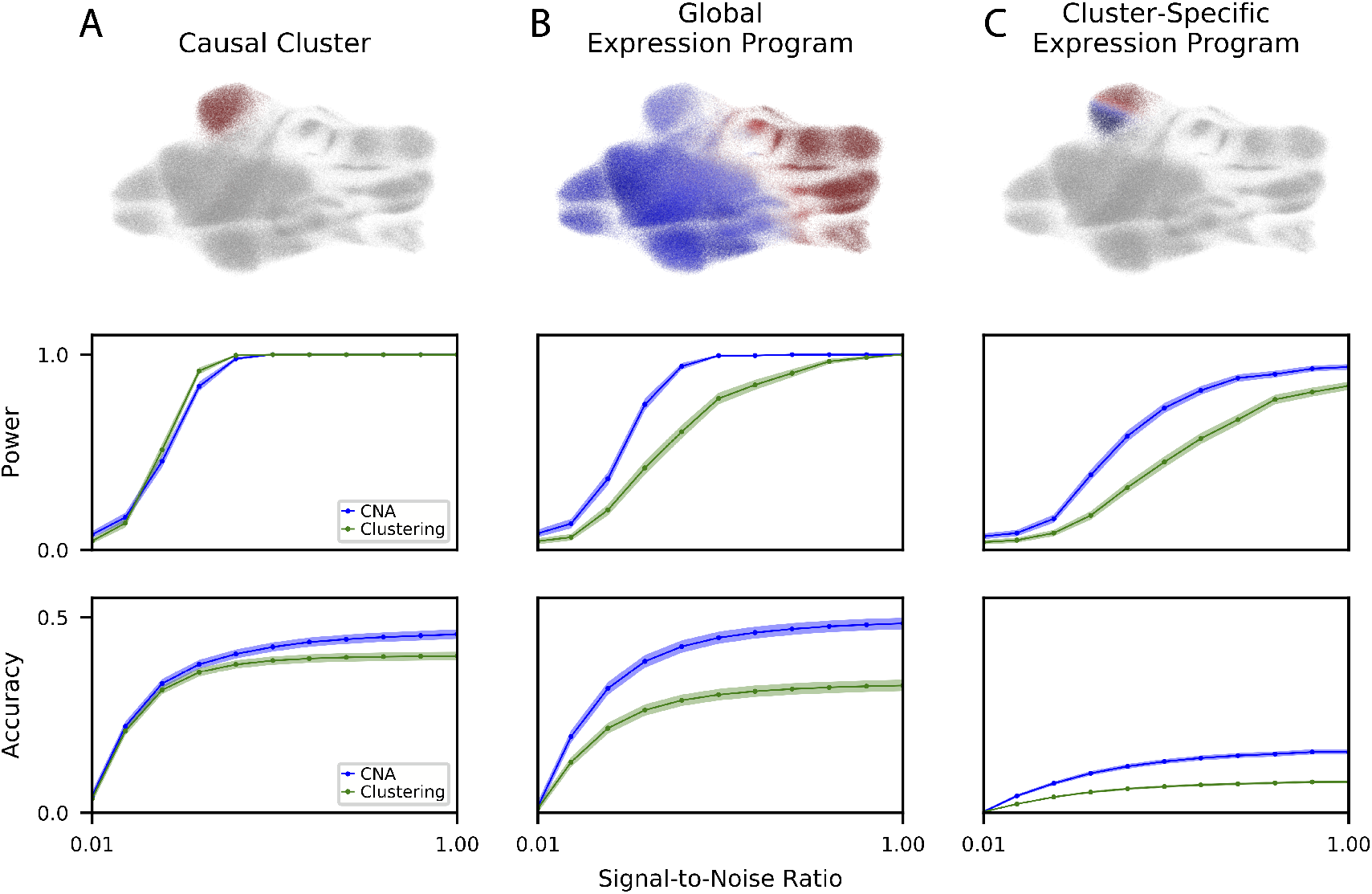
Power and accuracy assessed in simulation. We simulated three ground truth signal types: (**A**) causal clusters, (**B**) global gene expression programs, and (**C**) cluster-specific gene expression programs. For each signal type, we show (top) an example of the signal in UMAP space, (middle) the power of CNA versus a cluster-based approach (MASC) across a range of signal-to-noise ratios at ɑ = 0.05, and (bottom) the accuracy of CNA versus a cluster-based approach across a range of signal-to-noise ratios. For power and accuracy, we plot the mean across all simulations at the given signal-to-noise ratio, as well as the standard error around the mean.

CNA had superior power over cluster-based analysis to detect global GEP and cluster-specific GEP signals, while retaining comparable power for cluster abundance signals (**Figure 2A-C**). These conclusions also hold with respect to the best-performing individual clustering resolution: for the global GEP and cluster-specific GEP signals CNA had better power than cluster-based analysis at the best-performing resolution, and for cluster abundance signals CNA had comparable power to cluster-based analysis run on the best-performing clustering resolutions, including the ground-truth resolution used to define the clusters (**Supplementary Figure 4**).

CNA also had superior accuracy relative to cluster-based analysis for all three signal types (**Figure 2A-C**). Moreover, for global GEP signals, CNA’s accuracy was superior to accuracy for cluster-based analysis even at the best-performing clustering resolution (**Supplementary Figure 4**). For cluster abundance signals, the only resolution parameter choice that obtained superior accuracy to CNA was the one used to create the simulated cluster signals. For cluster-specific GEPs, the only resolution outperforming CNA was the finest resolution tested, which included 72 clusters; all other resolutions were less accurate, and two had accuracy near zero (**Supplementary Figure 4**).

### CNA captures Notch activation gradient implicated in rheumatoid arthritis

To assess whether CNA can detect important biological structure in real data, we applied CNA to 27,216 fibroblast scRNA-seq profiles from synovial joint tissue of six rheumatoid arthritis (RA) patients and six patients with osteoarthritis[8]. The original publication, also by our group, used trajectory analysis to uncover a fibroblast trajectory corresponding to endothelial Notch signaling and found expansion of Notch-activated fibroblasts in RA. This prior study also identified two fibroblast clusters—representing the lining versus sublining synovium regions—and demonstrated sublining fibroblast expansion in RA.

CNA identifies NAM-PC1 as the dominant signal in this dataset: NAM-PC 1 explains 39% of the variance in the NAM while no other NAM-PC explains more than 12%. NAM-PC1 reflects Notch activation: cells’ expression of *PRG4*—an established Notch-response gene in the synovial joint tissue[11]—was most strongly correlated with their anchored neighborhoods’ NAM-PC1 loadings (Pearson r=0.79, p<1e-10), followed by expression of *FN1* (Pearson r=0.71, p<1e-10), a signaling molecule shown to activate Notch[12]. Further, two Notch gene sets were significantly enriched among all gene correlations to NAM-PC1 (“Vilimas NOTCH1 targets up” and “Reactome signalling by NOTCH”, FDR=0.0073 and FDR=0.019, respectively). Moreover, NAM-PC1 has a stronger correlation than the published trajectory to the experimentally defined Notch activation score from the original paper (Spearman r=0.56 vs r=0.43, p<0.01 by bootstrapped permutation test, **Figure 3A-C**). CNA’s focus on inter-sample abundance covariance information was useful for uncovering this structure: PC1 from naive transcriptional PCA of the cells-by-genes expression matrix has a low correlation (Spearman r=0.22) with Notch activation (**Figure 3D**). Notably, NAM-PC1 detected the Notch activation signal without the parameter tuning required by trajectory analysis.

**Figure 3:**
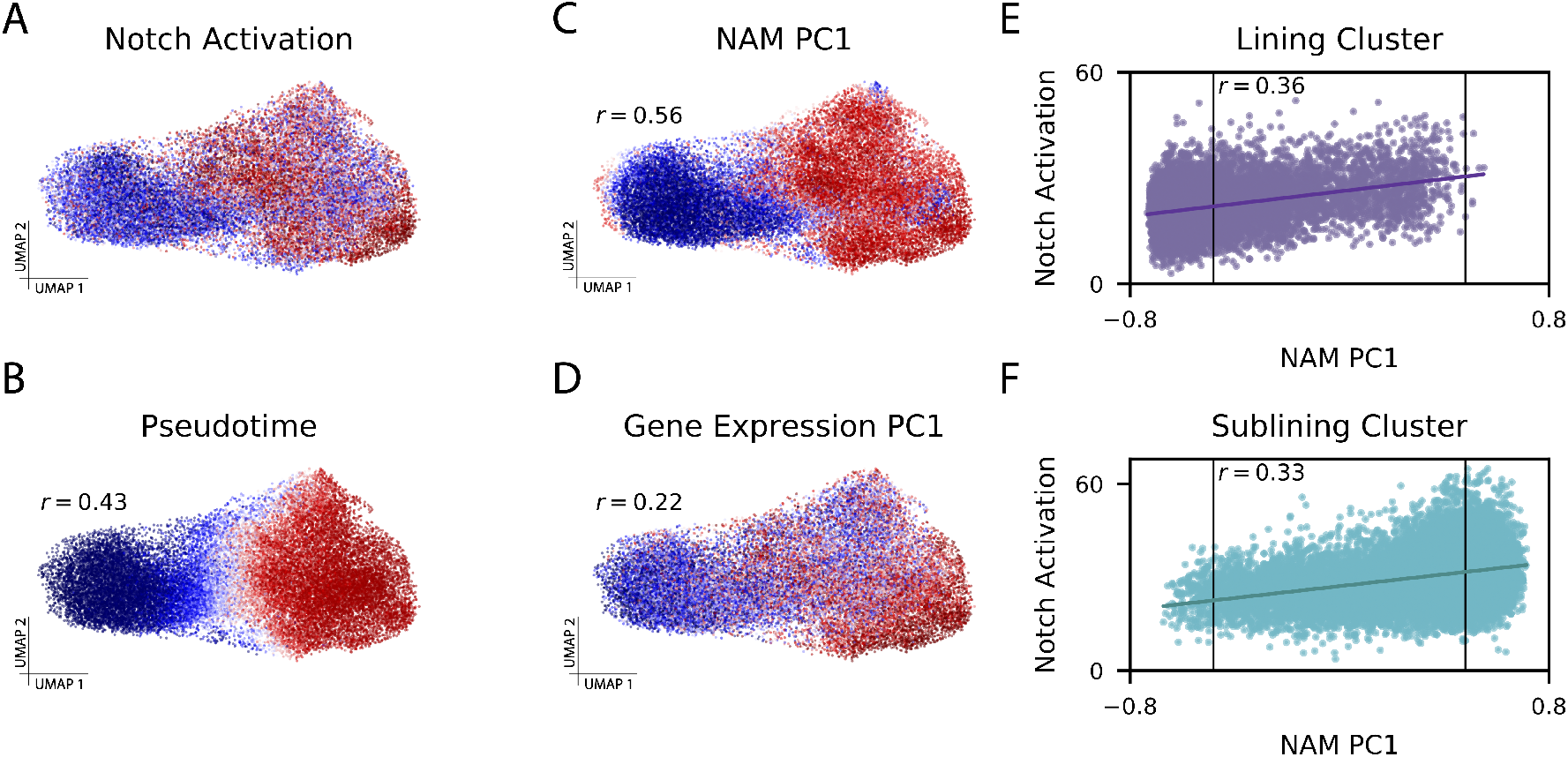
CNA captures Notch activation gradient in rheumatoid arthritis dataset. (**A**) Experimentally-determined Notch activation score per fibroblast cell. (**B**) Pseudotime assignments per cell (Spearman r=0.43 to Notch activation score). (**C**) The loading on NAM-PC1 for each cell’s anchored neighborhood (Spearman r=0.56 to Notch activation score). (**D**) Cell loadings on PC1 of the gene expression matrix (Spearman r=0.22 to Notch activation score). (**E**) Notch activation score per cell assigned to the lining cluster and (**F**) Notch activation score per cell assigned to the sublining cluster, each plotted against the anchored neighborhood’s loading on NAM-PC1. The FDR<0.05 thresholds beyond which each neighborhood was considered expanded in RA (right) or depleted in RA (left) are marked with vertical lines on (E) and (F), highlighting that some cells from each cluster are included in the expanded population and the depleted population.

NAM-PC1 largely separates the sublining and lining clusters (t-test p<1e-10) because sublining cells generally have higher Notch activation[8], but CNA reveals that Notch activation variation exists within these clusters. Neighborhood loadings on NAM-PC1 are correlated to the Notch activation scores of their anchor cells even within each cluster (Pearson r=0.36 lining cluster with p<0.001, Pearson r=0.33 sublining cluster with p<0.001; **Figure 3E-F**).

CNA identified RA-associated cell populations (global p=0.02) that recapitulate the coarse cluster-based associations but more precisely reflect the driving Notch mechanism. Nearly all cells to which CNA assigned significantly positive neighborhood coefficients (99.9% of 5,181 total cells at FDR<0.05) belong to the sublining cluster, and nearly all cells assigned significantly negative neighborhood coefficients belong to the lining cluster (96.8% of 7,169 total cells at FDR<0.05). However, CNA assigned some sublining-cluster cells to the depleted population, and these cells have lower Notch activation gene expression than other sublining-cluster cells. Likewise, CNA assigned some lining-cluster cells with higher Notch activation to the expanded population (**Figure 3E-F**). Therefore, CNA adds informative granularity beyond the cluster-based associations.

### CNA refines sepsis-associated blood cell populations

To assess CNA’s ability to identify granular case-control associations in a dataset with many cell types, we next applied CNA to scRNA-seq profiles of 102,814 peripheral blood mononuclear cells (PBMCs) from 29 patient with sepsis and 36 patients without sepsis. The published analysis[7] compared patients with and without sepsis in several sub-cohorts, *e*.*g*., among intensive care patients and among emergency department patients. Using a clustering of the data (**Figure 4A**), this analysis identified expansion of a monocyte state “MS1” in sepsis in multiple sub-cohorts (**Supplementary Table 2**). Our re-analysis compares patients with and without sepsis across the full cohort using CNA and a MASC cluster-based analysis.

**Figure 4:**
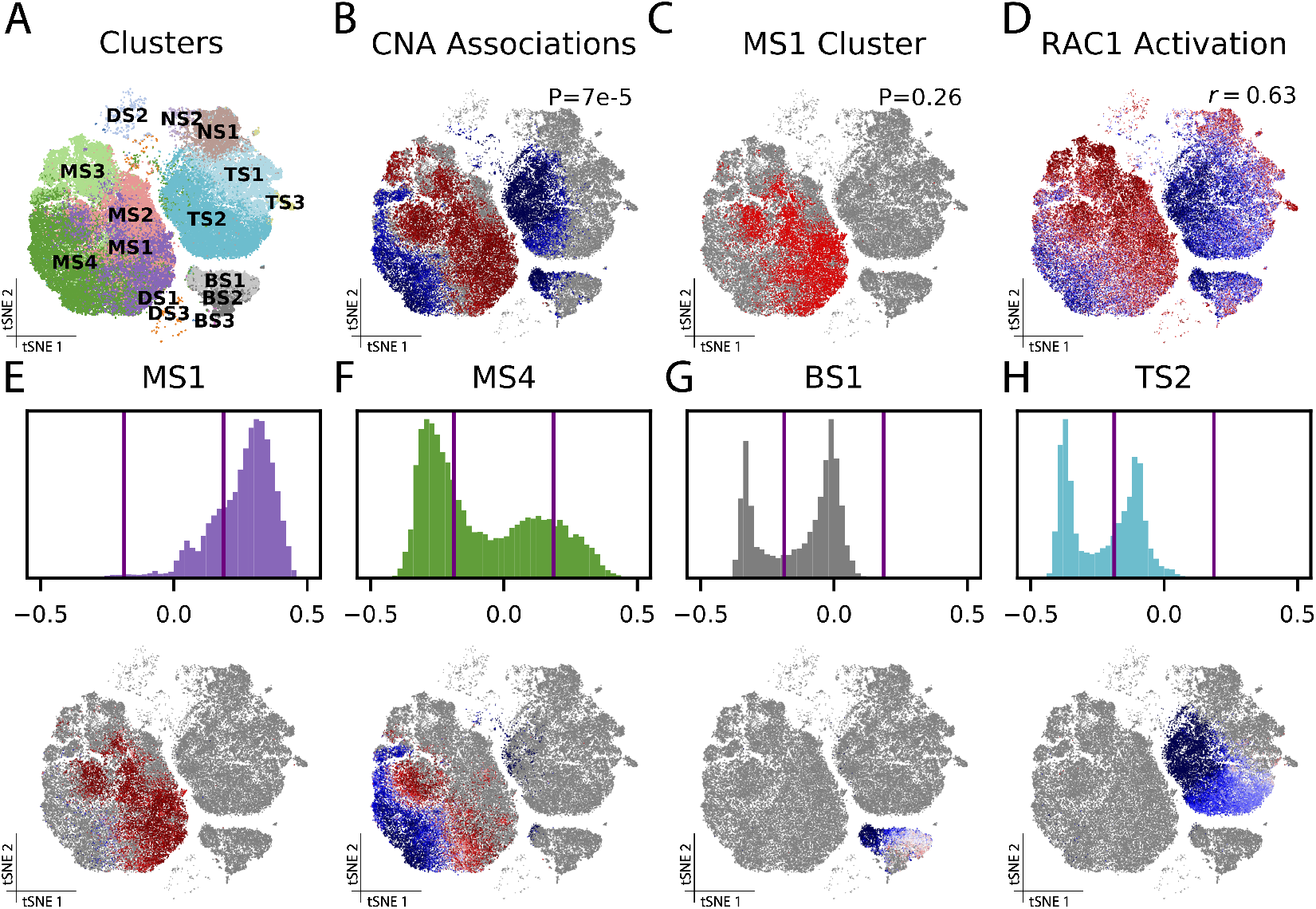
CNA refines sepsis-associated blood cell populations. (**A**) Clusters from the published analysis. (**B**) Results of association test for sepsis across the whole cohort using can (p=7e-5): each cell is colored according to its neighborhood coefficient, with red indicating high correlation and blue indicating low correlation. (**C**) MS1, the cluster closest to approaching nominal significance (p=0.26) in a cluster-based association test for sepsis across the whole cohort. (**D**) Cells colored according to their summed expression of genes in the RAC1 activation gene set. (**E-H**) The distribution of neighborhood coefficients to sepsis phenotype within several of the original clusters—MS1 (**E**), MS4 (**F**), BS1 (**G**) and TS2 (**H**)—are shown as histograms (top) and in tSNE space (bottom).

CNA found significant changes in sepsis compared to control samples (global p=7e-5) and identified a population expanded in sepsis (19,991 monocytes at FDR<0.05; **Figure 4B**). This population overlapped with MS1 but contained cells from other clusters: 56% of cells in CNA’s expanded population were in MS1 while 44% were in clusters MS2, MS3, and MS4. CNA’s expanded population contained 75% of all MS1 cells. In contrast, our cluster-based analysis of the same sepsis phenotype found that no cluster was significantly associated, though MS1 did have the smallest p-value (p=0.26; **Figure 4C**). Therefore, our results support the original finding but demonstrate that the published clusters partition transcriptional space in a manner that reduces power to detect the sepsis association in the full cohort.

CNA’s cluster-free delineation of sepsis-associated cell states implicates known sepsis-relevant pathways. Gene expression correlations to per-cell neighborhood coefficients were most highly enriched for the RAC1 activation gene set (FDR=2.5e-4, r=0.63 between summed gene-set expression and neighborhood coefficients), a known sepsis-associated pathway[13] whose suppression has therapeutic benefit in septic encephalopathy[14] (**Figure 4D**). The other most significantly enriched gene sets also have established sepsis associations (**Supplementary Table 3**).

Strikingly, CNA revealed considerable within-cluster heterogeneity in this dataset: eight of the fifteen published clusters included clear subpopulations with distinct degrees—and even directions—of associations to sepsis (**Figure 4E-H; Supplementary Figure 5**). For example, MS4 contains both a significantly expanded and a significantly depleted subpopulation (FDR<0.05; **Figure 4F**). Both of these associations were obscured by aggregating these subpopulations together. In the published analysis, clustering resolution was tailored to each cell type (e.g. Leiden 0.6 for T cells, 0.4 for monocytes). In contrast, CNA does not require parameter tuning to detect associated populations.

For comparison, we also ran MELD on this dataset. MELD per-cell abundance relationship scores to sepsis were correlated with CNA’s neighborhood coefficient values (r=0.6, **Supplementary Figure 6**). In contrast to CNA, however, MELD does not assess significance for these scores and produced patterns of scores on randomly permuted case-control labels that also appear to have nontrivial structure (**Supplementary Figure 6**). When we applied a permutation-based approach identical to the one used by CNA to assess significance at the neighborhood level, none of the individual per-cell MELD scores were significant at FDR<0.05 (**Supplementary Figure 6**). This highlights the power advantage of CNA’s use of inter-sample covariance information.

### CNA captures diverse associations in tuberculosis dataset

We next applied CNA to a larger and more richly phenotyped dataset: 500,089 memory T cells from 259 patients in a tuberculosis progression cohort[6]. (This dataset, recently published by our group, was also used above for simulations.) The published analysis employed 31 clusters to compare patients previously infected with *Mycobacterium tuberculosis* who rapidly developed symptoms (‘progressors’, N=128) to those who sustained latent infections (‘non-progressors’, N=131).

The NAM-PCs in this dataset appear to carry biologic meaning. For example, neighborhood loadings on NAM-PC1 correlate strongly across cells with a previously-defined transcriptional signature of “innateness,” the degree of effector function in each cell (r=0.81; **Figure 5A**)[15, 16], and individual gene correlations to NAM-PC1 also reflect this (**Supplementary Table 4**). This result shows that individuals vary to a substantial degree in their average T cell “innateness.” Moreover, NAM-PC1 sample loadings were nearly identical when computed using protein profiling, mRNA profiling, or the joint CCA representation of this data (**Figure 5B**); by contrast, PC1 sample loadings from naive PCA of each data type were far less correlated (**Figure 5B**). Across the three modalities, 50% of total variance in each NAM was explained by the top 5-10 PCs (out of 271; **Figure 5C**), suggesting NAM-PCs offer a parsimonious representation of this dataset.

**Figure 5:**
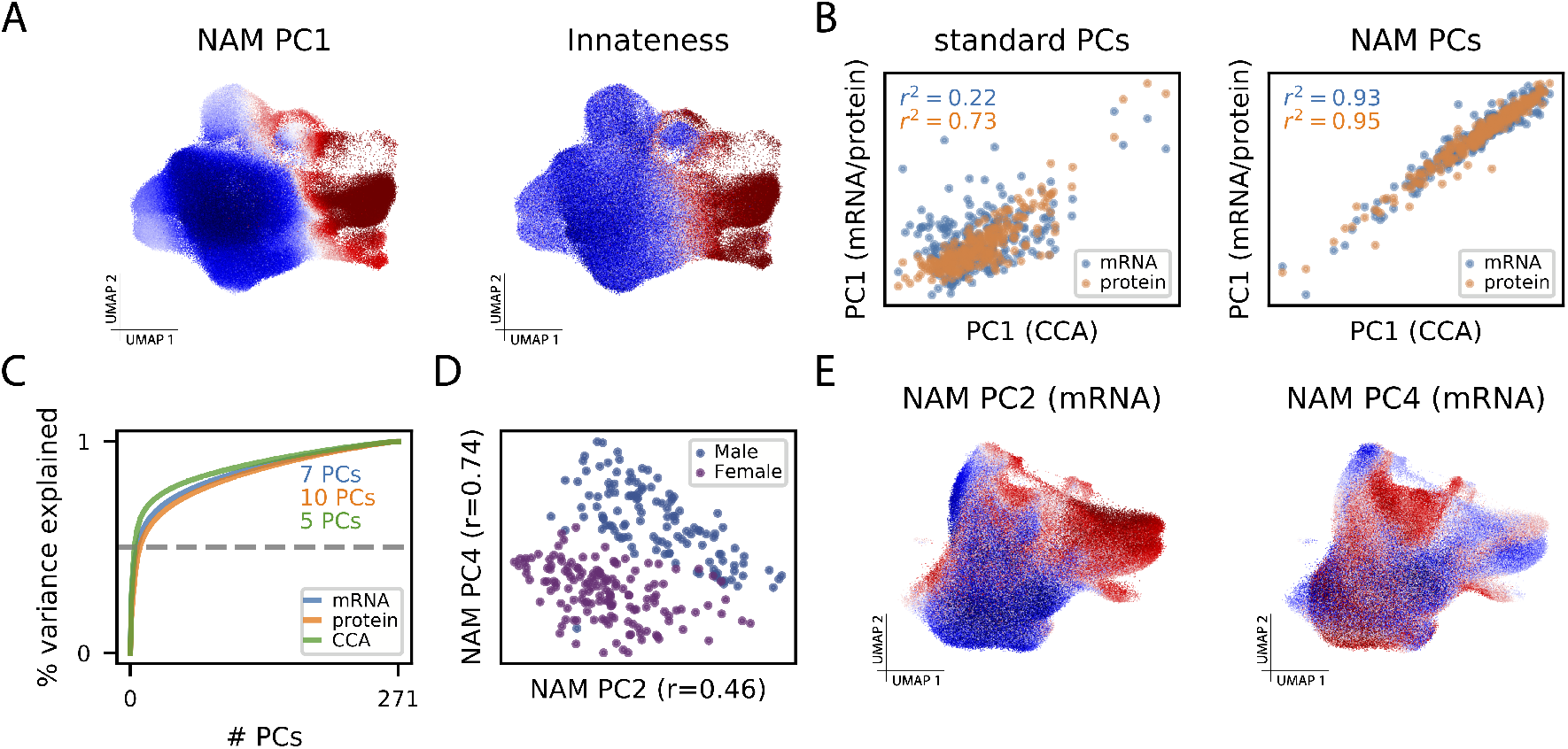
CNA characterizes biologically meaningful structure in TB dataset. (**A**) UMAP with cells colored by their anchored neighborhood’s loading on NAM-PC1 (left) or the transcriptional score of innateness from Gutierrez-Arcelus, *et al*. (right)[15]. (**B**) (Left) Sample loadings along the first PCs resulting from naive PCA of mRNA expression and protein expression plotted against the same loadings for the CCA-based joint mRNA/protein representation. (Right) Sample loadings along the first PCs resulting from PCA of the NAM generated from mRNA expression and protein expression plotted against the same loadings for the CCA-based joint mRNA/protein representation. (**C**) The cumulative percent of variance in the NAM explained by the NAM-PCs in each data modality. The minimum number of NAM-PCs needed to capture 50% of variance in each modality are highlighted. (**D**) Plot of sample loadings on NAM-PC2 and -PC4 colored by biological sex. (**E**) UMAP with cells colored by their anchored neighborhood’s loading on NAM-PC2 (left) or NAM-PC4 (right).

To assess whether NAM-PCs can detect nuanced transcriptional shifts that span a broad range of cell types, we re-computed NAM-PCs for this dataset without the upstream batch correction from the published analysis, which has the potential to eliminate subtle biologic variation[17, 18]. Indeed, in the mRNA data we found that NAM-PC2 and -PC4 correlate strongly with sex (joint R^2^=0.76; **Figure 5D**). Neighborhood loadings on NAM-PC4 indeed capture sex chromosome gene expression (**Supplementary Table 5**)—which differentiates otherwise very similar cells from individuals with different sex chromosomes across all cell types—while NAM-PC2 captures cell states known to vary in abundance with sex[19] (**Supplementary Table 6, Figure 5E**).

We next analyzed the primary phenotype, TB progression, using identical data processing and covariate control to the published analysis (**Methods**), which defined 31 clusters (**Figure 6A**) and found two clusters with Th17- and innate-like character (“C-12” and “C-20”, respectively) to be depleted among progressors. CNA found a significant global association (CNA global p=0.0015) driven by a depleted population (FDR<0.05) as well as an expanded population (FDR<0.05). CNA’s depleted population overlapped with the previously published C-12 (86% of cluster) and C-20 (64% of cluster) but also contained many cells from additional, phenotypically-similar clusters (74% of depleted population; **Figure 6B**). Overall, this population had similar characteristic proteins and genes to the cluster-based depleted population (**Figure 6C, Supplementary Table 7**) but contained substantially more cells.

**Figure 6:**
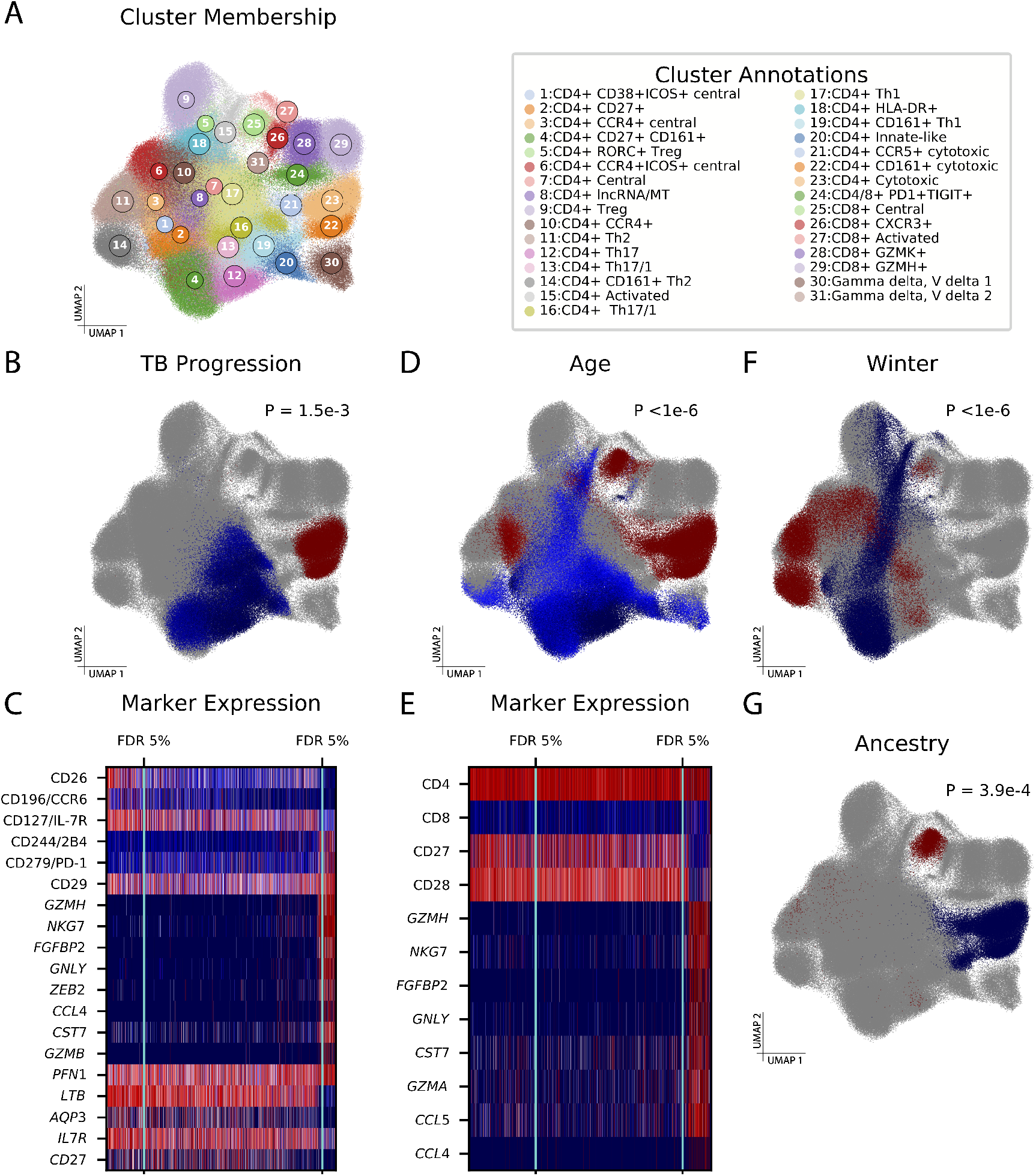
CNA improves characterization of diverse sample attributes in a tuberculosis cohort. (**A**) UMAP of memory T cells colored by cluster assignment in the original study. (**B**) Cells in UMAP space, colored according to their neighborhood’s abundance correlation to TB progressor versus non-progressor status, with red indicating high correlation and blue indicating low correlation. Cells whose neighborhood coefficients did not pass a FDR<0.05 threshold for association are shown in grey. **(C)** A heatmap of expression for genes and proteins of biological interest across cells, with red indicating high expression and blue indicating low expression. The cells are ordered from left to right according to their neighborhood coefficients to TB progressor status, and the FDR <0.05 thresholds beyond which cells are included in the significantly depleted (left) or expanded (right) population are shown in aqua. (**D**) Populations expanded and depleted with increasing age. **(E)** Heatmap of expression for genes and proteins of biological interest for the age association. (**F**) Populations expanded and depleted among samples drawn during the winter season relative to samples drawn during other seasons. (**F**) Populations expanded and depleted with increasing global fraction of European genetic ancestry.

In contrast to the cluster-based analysis, CNA newly revealed a population of cytotoxic cells expanded among progressors (**Figure 6B-C; Supplementary Table 7**), consistent with prior work describing the interplay between cytotoxic cells and mycobacteria[20]. These cells were predominantly captured by two clusters: 72% were from cluster C-23 (“CD4+ cytotoxic”) and 27% were from C-22 (“CD4+ CD161+ cytotoxic”). Tested individually, these clusters show weak evidence of association with progressor status (p=0.013 and 0.022, respectively) and do not pass multiple testing correction for the 31 clusters total. With a single test, CNA detected an associated population of functionally similar cells that had been split across multiple clusters.

In the original publication, the association to progressor status was only significant after unbiased mRNA profiling was combined with targeted surface protein quantification in a multimodal representation. Given our observed correlations among NAM-PCs across data modalities, we speculated that CNA might identify this association in unbiased mRNA data alone, and indeed it does (global p=4.5e-3).

Finally, we conducted a survey for associations between the single-cell data (multimodal representation) and 17 sample-level attributes besides progressor status (**Methods**). With control for confounders and multiple testing (**Methods; Supplementary Table 8**), we found global associations for age (p<1e-6; **Figure 6D-E**), season of blood draw (p<1e-6; **Figure 6F**), genetic ancestry (p=1.8e-4; **Figure 6G**), and sex (p=3e-6) (**Supplementary Figure 7, Supplementary Table 9**). These results align with the published cluster-based analysis, which also found evidence of these associations, and demonstrate that CNA can detect associations to a variety of signals, including demographic, environmental, and genetic factors. On average, CNA chose just 19 (out of 271; 7%) NAM-PCs to explain 26% of variance in these attributes, a 3.7x enrichment, suggesting that NAM-PCs are a parsimonious, phenotype-relevant representation of this complex dataset.

Some of the distinguishing cell states CNA finds associated with these attributes are established in the literature, while others are less well elucidated. We find older age is associated with higher CD8+/CD4+ ratio, decreased costimulatory molecule expression, and greater effector memory character relative to central memory character [21] (**Figure 6E; Supplementary Table 10)**. CNA highlights a shift toward more Th2 character relative to Th1 character during the winter season in contrast to studies of seasonality in other locations [22], and identifies an expanded CD8+ central memory population and depleted CD4+ cytotoxic population with increasing European genetic ancestry (**Supplementary Tables 11 and 12**). Cluster-based analyses with identical covariate control produced generally similar results, but implicated fewer cells in each association than CNA (**Supplementary Figure 7, Supplementary Table 9**).

## Discussion

In this work we introduced CNA, a method to characterize dominant axes of abundance variation across samples in a single-cell dataset and to identify with greater flexibility and granularity cell populations whose abundance correlates with sample attributes of interest. CNA offers improved power and accuracy over traditional cluster-based analysis while remaining robust to experimental artifacts and providing control for sample-level confounders, and it does so without requiring parameter tuning or long computation times. CNA can be used to study diverse sample attributes, enabling improved understanding of disease pathology, risk, and treatment.

In addition to their utility for testing for associations to sample-level attributes, NAM-PCs themselves appear to carry biological meaning: for example, our analyses revealed NAM-PCs that correspond to Notch signaling, memory T cell innateness, and a sex chromosome gene signature. For NAM-PCs without clear biological interpretation, characterizing the cellular functions and/or regulatory influences that unify these covarying neighborhood groups could yield insight into basic biology. Covarying neighborhood groups may, for example, delineate cell states most relevant to context-dependent cellular processes such as gene regulation and cellular metabolism.

CNA offers a versatile framework that can be easily extended to other data modalities. We highlight datasets of scRNA-seq and multimodal mRNA-and-protein profiling, but CNA can be extended to any modality for which cell-cell graphs can be built such as single-cell ATAC-seq epigenome profiling or mass cytometry protein profiling. For some of these applications, NAM factorization with approaches besides PCA, such as non-negative matrix factorization or independent components analysis, may be useful.

CNA has several limitations: first, CNA is likely to be less powerful than cluster-based methods on very small samples sizes (N<10). Second, though existing approaches for biological annotation of clusters and trajectories can be applied to CNA populations and NAM-PCs, respectively, such approaches typically seek a single explanatory signal; an associated population or NAM-PC might capture multiple related processes, and a given biological process may be captured by multiple NAM-PCs. Third, while cell assignments into clusters are discrete, CNA’s neighborhoods have probabilistic distributions in transcriptional space. As a result, it is not always obvious where the boundary of a CNA-associated population lies, or whether such a boundary exists.

Despite these limitations, CNA is a powerful new way to identify disease states and drivers of variation across samples in single-cell datasets that is unique in taking advantage of inter-sample variation. As single-cell datasets grow in sample volume, methods that leverage and characterize inter-sample information at fine-scale transcriptional resolution will become crucial to realize the promise of single-cell technologies.

## Acknowledgements

We thank Anika Gupta, Dylan Kotliar, Yang Luo, Nghia Millard, Miguel Reyes, Saori Sakaue, Fan Zhang, the members of the CGTA discussion group, and the Raychaudhuri lab for helpful discussions and feedback. This work is supported in part by funding from the National Institutes of Health (UH2AR067677, U19 AI111224, U01 HG009379, and 1R01AR063759). S.A. was supported by the Swiss National Science Foundation (SNSF) postdoctoral mobility fellowships P2ELP3_172101 and P400PB_183823 and NIH T32 grant T32HG010464.

## URLs

Open-source repository containing code for CNA: https://github.com/yakirr/cna

Open-source repository containing code underlying all figures and tables: https://github.com/yakirr/cna-display

## Methods

### Covarying Neighborhood Analysis

#### Intuition

Covarying neighborhood analysis is built on the idea of a *transcriptional neighborhood*, which is a very small subset of transcriptional space, typically much smaller than would arise from traditional clustering. Our method leverages two intuitions about transcriptional neighborhoods. First, because of the granularity of transcriptional neighborhoods, any meaningful variation across samples in a single-cell dataset will result in differential abundance of one or more neighborhoods across samples. Second, neighborhoods covary in abundance across samples because of shared function and/or regulatory influences. The first intuition leads us to represent multi-sample single-cell data using the *neighborhood abundance matrix* (NAM), a matrix of samples by neighborhoods that describes the relative abundance of each neighborhood in each sample. The second intuition leads us to analyze the NAM using principal components analysis. The resulting principal components tell us about both sets of neighborhoods that covary in abundance across samples as well as samples with similar abundance profiles across neighborhoods. This information is what we use to find structure and to conduct association testing with sample-level attributes such as clinical information, genotypic information, or experimental condition. The remainder of our technical material establishes notation and assumptions and then provides detailed descriptions of (i) definition of transcriptional neighborhoods, (ii) construction and quality control of the NAM, (iii) PCA of the NAM controlling for batch and covariates, and (iv) association testing.

#### Notation and assumptions

Let *X* be an *M* × *G=* matrix representing a single-cell dataset with *M* cells and *G* cell-level features such as genes. Let *N* be the number of distinct samples to which the cells belong, and for every cell *m* and every sample *n* let *C*(*n*) be the set of cells belonging to the *n*-th sample. We assume that *X* has already undergone quality control and (if desired) batch correction and that a nearest neighbor graph construction algorithm—such as the UMAP nearest neighbor algorithm—has been run on *X* to produce a sparse, weighted *M* × *M* adjacency matrix *A* whose *m, m*′-th entry indicates the similarity between cells *m* and *m*′ in the graph.

#### Definition of transcriptional neighborhoods

For each cell in our dataset, we define a transcriptional neighborhood anchored at that cell using the sense of locality provided by the nearest neighbor graph. That is, two cells can be considered to be close to each other if it is “easy” to reach one from the other in the graph. A natural way to define neighborhoods through this lens is to stipulate that two cells are in the same neighborhood to the extent that a random walk on the graph would be likely to reach one from the other.

More formally, we define a random walk whose transition probabilities are proportional to the entries of *I* + *A*, where *I* is the *M* × *M* identity matrix. (The addition of the identity adds self-loops to the UMAP graph.) That is, the probability that the walk moves from cell *m*′ to cell *m* in one step is given by

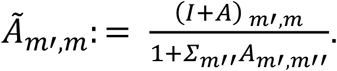

For some number of steps *s*, we then define the extent to which the *m*′-th cell belongs in the neighborhood of the *m*-th cell as the probability that a random walk starting at the *m*′-th cell will end up at the *m*-th cell after *s* steps. This is given by

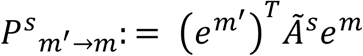

where *Ã* is the matrix whose entries are given by *Ã* _*m*′,*m*_ and *e*^*m*^ is a length-*M* vector whose *m*-th entry equals one and whose other entries are all zero, and *e*^*m*′^ is similarly defined. As we discuss in detail below, the number of steps *s* is chosen to minimize neighborhood size while ensuring that neighborhoods are not dominated by a small number of samples.

#### Construction and quality control of the neighborhood abundance matrix

With neighborhoods defined, we can now transform our dataset into a matrix of samples by neighborhoods whose *n, m*-th entry is the relative abundance of neighborhood *m* in sample *n*, i.e., the NAM. To formally define the NAM, we first let

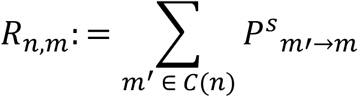

be the length-*M* vector representing the total number of expected cells from the *n*-th sample that would arrive in the *m*-th neighborhood as a result of our random walk. The NAM is then given by normalizing the rows of *R* to sum to one, i.e.,

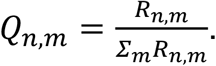

These entries can be computed very quickly using iterative sparse matrix multiplication, taking under one minute for a dataset with 500K cells and over 250 samples. See the Supplementary Note for details on efficient computation.

##### Choosing the length of the random walk

With the NAM defined, we can now discuss our choice of the number of steps *s* in the random walk that defines the NAM. Our guiding principle is that *s* should be chosen in a data-dependent manner to minimize neighborhood size, thereby retaining informative granularity, while ensuring that neighborhoods are not dominated by cells from a small number of samples. We quantify this by measuring, for each neighborhood, the kurtosis of its respective column of the NAM: a large kurtosis indicates that a small number of samples dominates the relevant neighborhood. With increasing timesteps, as the neighborhoods expand to incorporate more cells, kurtosis decreases. To achieve an appropriate balance between minimizing neighborhood size and eliminating unbalanced representation of samples in neighborhoods, we allow our random walk to continue until either the median kurtosis across neighborhoods is less than 8 (the kurtosis of a uniform distribution over only 10% of samples) or the median kurtosis across neighborhoods decreases by less than 3 (the kurtosis of the normal distribution) over consecutive time steps.

##### Removing neighborhoods with strong batch effects

If batch information is available, we also filter out neighborhoods that are dominated by one or a small number of batches by averaging the rows of the NAM within each batch to produce a batches-by-neighborhoods matrix and then computing for each neighborhood the kurtosis of its respective column of this new matrix. We then discard all neighborhoods with kurtosis greater than twice the median value across all neighborhoods.

##### Conditioning on sample-level covariates

If there is a set of sample-level covariates whose influence on *X* we do not wish to be represented among the principal components of the NAM, we linearly project them out of each column of the NAM, i.e., we regress each column of the NAM on the sample-level covariates that are supplied and replace it with the residuals arising from that regression. If there is batch information available, this can also be done with a one-hot encoding of batch IDs to further remove subtle batch effects that may still be present. In this case, we use ridge regression with an automatically chosen ridge parameter to account for the typically large number of batches relative to samples.

#### PCA of the NAM while conditioning on batch and covariates

Once the NAM is constructed, principal components analysis yields the decomposition

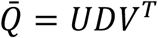

where 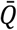 is the NAM with columns standardized to have mean zero and variance one; *U* is a matrix whose *i*-th column contains the *i*-th left singular vector, which has one entry per sample; *D* is the diagonal matrix of singular values; and *V* is a matrix whose *i*-th column contains the *i*-th right singular vector, which has one entry per neighborhood. Each of the right singular vectors identifies neighborhoods that covary in abundance across samples, and each of the left singular vectors identifies samples that have similar neighborhood abundance profiles across neighborhoods.

#### Association testing

CNA quantifies association of the NAM to a given sample-level attribute in two ways: i) a global quantification of the fraction of variance in the attribute explained by the single-cell data, with an associated p-value, and ii) a local estimate of the correlation between the attribute and each neighborhood’s abundance across samples, with associated false discovery rates.

##### Global association test

Let *y* be a length-*N* vector containing a sample-level attribute of interest such as clinical information, genotypic information, or experimental condition, and suppose we want to associate *y* with the inter-sample variation in *X*. Because the left left singular vectors of the NAM, i.e., the columns of *U*, each contain one number per sample, we can do this in a simple linear model in which each sample is an observation. That is, for some number *k* of principal components, we can fit the model

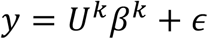

where *U*^*k*^ denotes the first *k* columns of *U, β*^*k*^ is a length-*k* vector with one coefficient per principal component, and *ϵ* represents mean-zero noise.

To choose *k* in a flexible and automatic way, we fit the above model for four different values of *k* ranging from [*N* /50] through *min*([*N* /50], *N* /5), where […] denotes the ceiling function. For each of these values of *k*, we compute a multivariate F-test p-value for the null hypothesis *H*_0_: *β*^*k*^ = 0, and we choose the value *k*^*^ that yields the minimal p-value. This ensures that larger values of *k* are only selected if they provide increased predictive power for *y* over and above what we would expect simply from their providing more degrees of freedom to the model.

If there are covariates that are residualized out of the NAM, these are residualized out of *y* prior to fitting the model. Similarly, if batch information has been residualized out of the NAM, it is likewise residualized out of *y* prior to fitting the model using the same ridge parameter used for the residualization out of the NAM.

To obtain a p-value for global association, we then perform the above procedure, including the selection of *k*, on a large number of empirical null instantiations (1,000 by default) obtained by permuting the values of *y* within each batch of the dataset. We then use the resulting set of p-values, of which there is one per null instantiation, as our null distribution.

##### Local association test

A natural notion of neighborhood-level effect size would be the correlation between the *m*-th column of the NAM and *y*. However, the NAM is very high-dimensional and so these correlations are noisy. Instead, we therefore compute a “smoothed correlation” between neighborhood *m* and *y*, i.e., the correlation between the *m*-th column of the rank-*k*^*^ representation of the NAM and *y*. Mathematically, this equals

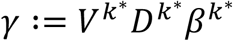

where *D*^*k*^ denotes the upper-left *k* × *k* submatrix of *D, V*^*k*^ denotes the first *k* columns of *V*, and *β*^*k*\*^ is the estimate from the model for global association. We refer to the entries of *γ* as the *neighborhood coefficients* for the sample attribute *y*. To assess statistical significance, we again compute null versions of *γ* and use these to estimate an empirical false discovery rate for a variety of magnitudes of correlation by comparing the number of entries of *γ* with magnitude above a given threshold to the average number of entries in the null versions of *γ* with magnitude above that threshold.

### Assessing performance with simulations

To assess the calibration and power of our method, we carried out simulations using real single-cell data from the tuberculosis research unit (TBRU) cohort[6]. This dataset consists of *M* = 500,089 memory T cells from *N* = 271 samples that were profiled with CITE-Seq[23], which simultaneously provides both single-cell RNA-seq and single-cell quantification of 31 surface proteins. (Further aspects of this dataset are described elsewhere in the Methods.) All of our simulations used a 20-dimensional canonical correlation analysis-based representation of each cell generated by the authors of the original study that incorporated information shared by the RNA-seq and surface protein modalities and that was subsequently run through the Harmony method[24] to remove potential sample-specific and batch effects. All simulations were conducted at the full sample size of N=271.

#### Quantifying type 1 error

For the simulation **in Supplementary Figure 2A**, we simulated 1,000 independent null attributes by permuting an existing sample-level covariate in the dataset (age at sample collection) across all samples in the dataset. For the simulation in **Supplementary Figure 2B**, we simulated 1,000 independent null attributes by permuting an existing sample-level covariate in the dataset (age at sample collection) within each batch. This was done to preserve whatever batch effects may be present in the data in our null attributes. For the simulation in **Supplementary Figure 2C**, we created 1,000 independent null attributes with maximal batch effect by selecting 1,000 batches {*b*_1_,…, *b*_<1000_} randomly with replacement and setting the *i*-th attribute to equal one for all the samples in batch *b*_*i*_ and zero otherwise.

In all of the above simulations, we used CNA to obtain a p-value for association to the single-cell data for each attribute, accounting for possible batch effects. This yielded 1,000 p-values in each case.

#### Quantifying Type 2 error: Generation of simulated attributes

For **Figure 2**, we simulated three different signal types: cluster abundance (**Figure 2A**), global gene expression program **(Figure 2B**), and cluster-specific gene expression program (**Figure 2C**). In each case, we added gaussian noise to each simulated attribute to achieve signal-to-noise ratios of {0.01,0.1,0.2, …,0.9,1}.

For the cluster abundance signal type (**Figure 2A**), we clustered the single-cell data using the Leiden clustering algorithm[4] with the same resolution (2.0) used by the authors of the original TBRU study[6]. We then filtered out any clusters that did not have at least 10 samples with at least 50 cells each, and we filtered out any clusters whose abundance had correlation greater than 0.25 to membership in any batch. This reduced the number of clusters from 26 to 24. For each of the remaining 24 clusters, we computed the abundance of that cluster. For each of our 11 signal-to-noise ratios, we then simulated 10 independent attributes by summing this attribute with the appropriate amount of gaussian noise. This resulted in 24×10 = 240 attribute per noise level.

For the global gene expression program signal type (**Figure 2B**), we treated the 20 per-cell harmonized canonical variables as each representing the activity of a gene expression program. For each canonical variable, we computed the average value of that variable across all the cells in each sample. For each of our 11 signal-to-noise ratios, we then simulated 10 independent attributes by summing this attribute with the appropriate amount of gaussian noise. This resulted in 20×10 = 200 attributes per noise level.

For the cluster-specific gene expression program signal type (**Figure 2C**), we first clustered our data using the Leiden clustering algorithm with a resolution of 1.0 and filtered the clusters using the same criteria as in cluster abundance simulation. (We used a coarser clustering resolution here because this signal type is driven by intra-cluster variability rather than inter-cluster variability, so we wanted to achieve larger clusters.) For each of the 10 largest clusters, we then computed the top 3 principal components of all the harmonized canonical variables among only the cells in that cluster, which we treated as each representing activity of a cluster-specific gene expression program. For each of these principal components, we computed the average value of that component across all the cells in each sample, assigning cells outside the cluster in question a score of zero. For each of our 11 signal-to-noise ratios, we then simulated 10 independent attributes by summing this attribute with the appropriate amount of gaussian noise. This resulted in 10×3×10 = 300 attributes per noise level.

#### Analysis of simulated attributes

For each signal type, we analyzed the simulated attributes using i) CNA accounting for possible batch effects, and ii) MASC with the recommended inclusion of sample-level and batch-level random effects. MASC requires a set of clusters whose abundance it assesses for correlation with attribute, and so we ran 4 different versions of MASC using 4 different sets of clusters. These were created by running Leiden clustering on our data with resolution parameters of 0.2, 1, 2, and 5, resulting in 3, 15, 26, and 72 clusters, respectively.

To estimate power for a given signal type, we computed for each method and for each signal-to-noise ratio the fraction of tests in which the method reported a p-value less than 0.05. For CNA, we used the global p-value. For MASC, we used the lowest p-value for any individual cluster, multiplied by the total number of clusters to achieve multiple testing correction. We quantified uncertainty in our power estimates by computing empirical standard errors for our estimate of this mean. In **Figure 2**, we aggregated the 4 versions of MASC into one p-value by computing the minimum p-value across all 4 MASC clustering resolutions and Bonferroni correcting this for 4 tests. **Supplementary Figure 3** shows results at the level of individual MASC clustering resolutions.

To define a notion of accuracy for each method, we first utilized each method to obtain per-cell estimates of correlation to the attribute as follows. For CNA, we used the neighborhood coefficient for each cell, which is the per-neighborhood correlation to the attribute from the neighborhood for which that cell is the anchor. For MASC at a given clustering resolution, we assigned to each cell the signed effect size beta that MASC estimated for that cell’s parent cluster. We then defined ground truth per-cell scores for each signal type such that the noiseless version of each attribute would be obtained by averaging the per-cell scores of all the cells in each sample: for the cluster abundance signal type, we assigned a score of one to cells in the causal cluster and a score of zero to other cells; for the global gene expression program signal type, we used the per-cell values of the canonical variable in question; and for the cluster-specific gene expression program signal type, we used the per-cell values of the principal component in question, with a score of zero assigned to all cells outside the cluster.

To estimate accuracy for a given signal type, we then computed for each method and for each signal-to-noise ratio the average per-attribute correlation between the method’s reported per-cell scores and the ground truth per-cell scores for that attribute. We quantified uncertainty in this estimate by computing empirical standard errors for our estimate of this mean. In **Figure 2**, we aggregated the 4 versions of MASC into one p-value by averaging their accuracies. **Supplementary Figure 3** shows results at the level of individual MASC clustering resolutions.

### Analyses of real data

We analyzed three real datasets: a dataset of synovial fibroblasts from patients with rheumatoid arthritis versus osteoarthritis (N=12 samples, M=27,216 cells)[8]; a dataset of peripheral blood mononuclear cells from patients with and without sepsis (N=65 samples, M=102,814 cells)[7]; and a dataset of memory T cells from patients in a large tuberculosis progression cohort (N=271 samples, M=500,089 cells)[6].

#### Analysis of rheumatoid arthritis dataset

We obtained the rheumatoid arthritis dataset from the authors of a study of synovium of patients with rheumatoid arthritis (RA) and osteoarthritis (OA). The dataset consisted of M=27,216 synovial cells from N=12 samples profiled with single-cell RNAseq and processed with the Harmony algorithm to mitigate sample-specific effects. The cells also had cluster labels, assigned by the authors of the original study, that segmented the fibroblasts into lining and sub-lining populations. Following the approach of the original study in computing trajectory analysis, we filtered this dataset to fibroblasts only. We then applied CNA to this dataset without covariate or batch correction to produce an NAM with principal components as well as a p-value for global association to RA/OA status and neighborhood-level correlations to RA/OA status with corresponding FDRs.

To assess whether NAM PC1 was related to Notch activation, we obtained the per-cell Notch activation scores defined using the experimentally derived Notch activation gene set from the original study and computed the correlation across cells between these scores and the NAM PC1 neighborhood loading assigned to each cell’s neighborhood. We computed a p-value for whether this was significantly greater than the corresponding correlation for the per-cell values provided by the original trajectory analysis by bootstrapping over samples to create a null distribution for the difference between the magnitudes of the two correlations.

To assess whether PC1 of the NAM was related to Notch activation at the gene-set level, we computed correlations for the top 5,000 most variable genes between expression level in each cell and the cells’ anchored neighborhood loadings on NAM-PC1. Using these per-gene correlations as our input ranked list, we computed the enrichment of gene sets containing the term “NOTCH” from MSigDB’s “C7” catalogue of immune-related gene sets. We used R’s FGSEA package[25] with a maximum gene set size of 500, a minimum size of 15, and 100,000 permutations.

#### Analysis of sepsis dataset

We downloaded the sepsis dataset of Reyes *et al*. from the Broad Institute Single Cell Portal. The dataset consisted of M=102,814 CD45+ peripheral blood mononuclear cells (PBMCs) from N=65 samples profiled with single-cell RNA-seq. The study consisted of three clinical cohorts: i) patients presenting to the emergency department (ED) with urinary tract infection, divided into patients with leukocytosis but no organ dysfunction (Leuk-UTI, N=10), urosepsis (Int-URO, N=7) and persistent urosepsis (URO, N=10); ii) bacteremic patients with sepsis in hospital wards (Bac-SEP, N=4); and iii) patients admitted to the intensive care unit (ICU) with either sepsis (ICU-SEP, N=8) or no sepsis (ICU-NoSEP, N=7). There were also 19 healthy controls (Control, N=19). In total, the study included 29 sepsis patients (Int-URO, URO, and Bac-SEP) and 36 non-sepsis patients (Control, Leuk-UTI, and ICU-NoSEP).

To maximize comparability between our analysis and the original analysis, we analyzed the data following the same preprocessing steps as the original authors, namely: we filtered out cells with fewer than 100 unique molecular identifiers (UMIs) and genes with expression in fewer than 10 cells, and then we log-normalized the counts and filtered out genes with mean expression less than 0.0125 or dispersion less than 0.5. The dataset also includes some samples that are enriched for dendritic cells; following the original analysis, we included these in the dataset but did not assign them phenotype labels so that they could be included in the unsupervised portion of the analysis but would not directly affect any of the association analyses.

The original publication conducted case-control comparisons within 9 different subgroups of sepsis patients and controls (e.g., {URO, Int-URO} vs {Control, Leuk-UTI}). We conducted the same 9 association tests using CNA (without batch information or covariates, following the original study) and found good qualitative agreement; see **Supplementary Table 5**.

We then ran CNA on the aggregate phenotype of “any sepsis”, for which sepsis was defined as {Int-URO, URO, Bac-SEP, ICU-SEP} and non-sepsis was defined as {Control, Leuk-UTI, ICU-NoSEP}. To assess for gene set enrichment in association with sepsis, we computed correlations for the top 5,000 most variable genes between expression level in each cell and the cells’ anchored neighborhood loadings on NAM-PC1. Using these per-gene correlations as our input ranked list, we computed the enrichment of gene sets from the Pathways Interaction Database (“PID”) stored in MSigDB’s “C2” catalogue of curated gene sets. We used R’s FGSEA package[25] with a maximum gene set size of 500, a minimum size of 15, and 100,000 permutations.

To assess for heterogeneity within the 15 author-defined cell states, we then examined the distribution of CNA-estimated neighborhood correlations to the “any sepsis” phenotype within each author-defined cell-state (see **Supplementary Figure 4**). We identified MS4, TS2, and BS1 as the most visually striking examples of bimodality of these correlations within individual clusters.

#### Analysis of tuberculosis dataset

We obtained the pre-processed TBRU dataset directly from the authors of the index study^7^. This dataset consisted of M=500,089 memory T cells from N=271 samples that were profiled with CITE-seq[23], which simultaneously provides both single-cell RNA-seq and single-cell quantification of 31 surface proteins. The TBRU cohort was designed to identify correlates of progression to active tuberculosis infection compared to latent tuberculosis infection. Accordingly, approximately half the samples come from patients who had active TB at enrollment (4 - 7 years before the single-cell data were collected) and half the samples come from household contacts of these patients who developed latent infections after enrollment. In addition to this information, the dataset also contains a variety of other sample-level attributes such as age, sex, weight, and ancestry imputed from genotype information about each sample. See Supplementary Table 6 for the full list of sample-level information that we analyzed with CNA.

The authors of the original study used canonical correlation analysis to create a 20-dimensional representation of each cell that incorporated information shared by the RNA-seq and surface protein modalities and that was subsequently run through the Harmony method[24] to remove potential sample-specific and batch effects. To maximize comparability between our analysis and the original analysis, we used this representation in all analyses unless stated otherwise.

##### Unsupervised analysis

We computed the initial NAM by running CNA on the full dataset with correction only for batch and per-sample averages of i) the percent mitochondrial reads (pMT) of each cell, and ii) the number of unique molecular identifiers (nUMI) for each cell. To identify biological processes corresponding to NAM PCs, we then computed the correlation per-gene (the top 5,000 most variable in the dataset) and per-protein between expression level in each cell and the cells’ anchored neighborhood loadings on each NAM-PC.

##### Association analysis for TB progression phenotype

We analyzed the TB progression attribute with CNA, controlling for the same covariates that the authors of the original study used in their analysis: pMT, nUMI, age, age squared, sex, season of blood draw, and percent European ancestry. We retained cells whose neighborhood coefficients showed correlation to TB progression at FDR < 0.05. These cells clearly segregated into two contiguous groups in UMAP space: a depleted population and an enriched population. We examined the genes, among the top 5,000 most variable, and the surface proteins whose expression per-cell was most highly correlated with the cells’ anchored neighborhoods’ estimated abundance correlations to the TB phenotype.

##### Association survey across many sample-level attributes

We one-hot encoded all categorical attributes and standardized all continuous attributes. We then filtered the attributes by removing any attributes with missing values for >10% of samples, by removing any one-hot categorical attributes with fewer than 20 individuals represented, and by removing one from every attribute pair with a correlation>0.75. 17 of the attributes were retained after this step. We then determined for each of these 17 attributes *y* which others (including TB progression status) had a nominally significant (p<0.05) correlation to *y* and included those as covariates when analyzing *y*. Using the resulting selected covariates, shown in **Supplementary Table 6**, we ran CNA. Using the 31 clusters previously identified in this data, we ran a per-cluster association test with identical covariate control for each attribute. We added multiple hypothesis testing correction across clusters, and, for both cluster-based analysis and CNA, across the 17 attributes tested. For each attribute with a globally-significant association by CNA, we examined the genes, among the top 5000 variable genes, and the surface proteins whose expression per anchor cell was most highly correlated with corresponding neighborhood coefficients to the given attribute.

## Data Availability

All data analyzed during this study were available through three previously-published articles [6-8].

## Code Availability

As reported under the **URLs** section, an open-source repository containing code for running CNA is available at https://github.com/yakirr/cna and an open-source repository containing code underlying all figures and tables is available at https://github.com/yakirr/cna-display.

