## Supplementary Figures and Tables for "Axes of inter-sample variability among transcriptional neighborhoods reveal disease-associated cell states in single-cell data"

#### Contents

**Supplementary Tables** **2**

**Supplementary Figures** **8**

### Supplementary Tables

| Dataset | # cells | # steps | Avg. nbhd. size |
|---|---|---|---|
| RA | 27216 | 3 | 336.43 |
| Sepsis | 102814 | 4 | 1503.33 |
| TB | 500089 | 7 | 12183.74 |

**Supplementary Table 1: Neighborhood characteristics for each dataset analyzed**. For each of the three datasets analyzed in this paper, we show the number of random walk steps selected by CNA as well as the average across neighborhoods of the number of cells required to capture 70% of the mass of each neighborhood, estimated by sampling 500 random neighborhoods.

| **Phenotype** | | **Reyes et al.** | **CNA** | | |
|---|---|---|---|---|---|
| Cases | Controls | MS1 FDR | Global P | # NAM-PCs | # nbhds with FDR< 10% |
| Int-URO, URO, Bac-SEP, ICU-SEP | Control, Leuk-UTI, ICU-NoSEP | NA | $7.0 \times 10^{-5}$ | 2 | $50,696$ |
| URO, Int-URO | Control, Leuk-UTI | $< 10^{-3}$ | $2.8 \times 10^{-4}$ | 2 | $25,875$ |
| Bac-SEP, ICU-SEP | Control | $< 10^{-3}$ | $5.0 \times 10^{-4}$ | 2 | 0 |
| URO, Int-URO Bac-SEP, ICU-SEP | ICU-NoSEP | 0.27 | 0.86 | 3 | 0 |
| Leuk-UTI | Control | $6.0 \times 10^{-2}$ | 0.21 | 4 | 0 |
| Int-URO | Control | $< 10^{-3}$ | $5.9 \times 10^{-4}$ | 2 | 0 |
| URO | Control | $< 10^{-3}$ | $1.5 \times 10^{-2}$ | 4 | 0 |
| Bac-SEP | Control | $3.0 \times 10^{-3}$ | 0.12 | 2 | 0 |
| ICU-SEP | Control | $< 10^{-3}$ | $2.6 \times 10^{-3}$ | 2 | 0 |
| ICU-NoSEP | Control | $< 10^{-3}$ | $1.6 \times 10^{-2}$ | 2 | 0 |

**Supplementary Table 2: Sub-cohort assessments for sepsis dataset**. The original authors did not perform an aggregated sepsis versus non-sepsis association test. Rather, they compared every patient group to healthy controls (6 tests), in addition to the following tests: Int-URO and URO vs. Control or Leuk-UTI, Bac-SEP and ICU-SEP vs. Control, Int-URO, URO, Bac-EP and ICU-SEP vs. ICU-NoSEP. We performed association tests for these same 9 patient groupings using CNA as well as an aggregated sepsis vs. no sepsis phenotype (Int-URO, URO, Bac-SEP, and ICU-SEP vs. Leuk-UTI and Controls). The aggregated phenotype analysis (sepsis vs. no sepsis) was used for downstream analysis.

| Pathway | Enrichment | P, Adjusted | NAM PC1 corr | NAM PC2 corr |
|---|---|---|---|---|
| RAC1 | 0.71 | 0.0003 | 0.42 | 0.47 |
| PDGFRB | 0.52 | 0.0017 | 0.52 | 0.39 |
| ERBB1_DOWNSTREAM | 0.54 | 0.0023 | 0.50 | 0.38 |
| TOLL_ENDOGENOUS | 0.74 | 0.0024 | 0.17 | 0.61 |
| CDC42 | 0.61 | 0.0029 | 0.44 | 0.40 |
| TXA | 0.64 | 0.0037 | 0.38 | 0.39 |
| IL6_7 | 0.58 | 0.0160 | 0.34 | 0.43 |
| IL8_CXCR2 | 0.62 | 0.0170 | 0.34 | 0.37 |
| AMB2_NEUTROPHILS | 0.65 | 0.0212 | 0.43 | 0.32 |
| LYSOPHOSPHOLIPID | 0.60 | 0.0257 | 0.32 | 0.45 |
| CASPASE | 0.56 | 0.0298 | 0.51 | 0.11 |
| P38_ALPHA_BETA | 0.63 | 0.0305 | 0.26 | 0.41 |
| THROMBIN_PAR1 | 0.59 | 0.0340 | 0.37 | 0.33 |
| EPO | 0.60 | 0.0409 | 0.32 | 0.29 |
| ECADHERIN_NASCENT_AJ | 0.58 | 0.0489 | 0.33 | 0.33 |

**Supplementary Table 3: Gene sets enriched among transcriptional regions with higher abundance among sepsis patients**. Gene-set enrichment analysis was performed where the input rank list of genes was computed based on the correlation across cells of gene expression per anchor cell and the corresponding neighborhood's estimated abundance correlation to the sepsis phenotype. Many of these gene sets have established links to sepsis: *PDGFRB* knockout attenuates brain inflammation in sepsis,[1] ERBB1 (EGFR) inhibition in vivo blocks septic shock,[2] TLR signaling contributes to cytokine storms in sepsis,[3] and *CDC42* is beneficially upregulated in sepsis to restore endothelial barrier function and decrease edema.[4]

| Name | Type | Correlation | P.value |
|---|---|---|---|
| CD62L | Protein | 0.15 | $<$1e-10 |
| SELL | Gene | 0.24 | $<$1e-10 |
| NKG7 | Gene | -0.81 | $<$1e-10 |
| GZMH | Gene | -0.76 | $<$1e-10 |
| CCL5 | Gene | -0.70 | $<$1e-10 |
| GZMA | Gene | -0.67 | $<$1e-10 |
| GNLY | Gene | -0.67 | $<$1e-10 |

**Supplementary Table 4: Gene expression correlations to neighborhood loading on NAM-PC1 reflect a spectrum of "innateness"**. Correlations were computed between per-cell gene expression and the cell's anchored neighborhood's loading on NAM-PC1 from the joint CCA representation of this dataset. Cells with higher transcriptional 'innateness' signature tended to have lower loadings on NAM-PC1. Consistent with this pattern, CD62L/*SELL* expression is positively correlated with neighborhood loading on NAM-PC1 and expression levels of effector molecules are negatively correlated with neighborhood loading on NAM-PC1.

| Name | Type | Correlation | P.value |
|---|---|---|---|
| XIST | Gene | -0.41 | <1e-10 |
| RPS4Y1 | Gene | 0.58 | <1e-10 |
| DDX3Y | Gene | 0.35 | <1e-10 |
| UTY | Gene | 0.30 | <1e-10 |
| TTTY15 | Gene | 0.26 | <1e-10 |
| KDM5D | Gene | 0.18 | <1e-10 |
| USP9Y | Gene | 0.16 | <1e-10 |

**Supplementary Table 5: Sex chromosome genes are the most correlated in expression with NAM-PC4**. Correlations between per-cell gene expression and cell loading on NAM-PC4 were computed. The genes whose expression was most correlated or most anti-correlated with cell loading, shown here, were located on sex chromosomes. For each of these genes, a simple linear model was used to evaluate significance for the association between expression level and cell loading on NAM-PC4.

| Name | Type | Correlation | P.value |
|---|---|---|---|
| CD4 | Protein | -0.27 | <1e-10 |
| CD8 | Protein | 0.33 | <1e-10 |

**Supplementary Table 6: Differential expression of surface proteins on NAM-PC2 reflect cell population changes associated with sex chromosome differences**. Differential expression analysis was performed where the input rank list of genes was computed based on the correlation across cells between gene expression and cell loadings along NAM-PC2 of the mRNA profiling, without upstream batch correction from the original publication.

| Name | Type | Correlation | P.value |
|---|---|---|---|
| CD26 | Protein | -0.39 | <1e-10 |
| CD196/CCR6 | Protein | -0.41 | <1e-10 |
| CD127/IL-7R | Protein | -0.28 | <1e-10 |
| CD244/2B4 | Protein | 0.44 | <1e-10 |
| CD279/PD-1 | Protein | 0.20 | <1e-10 |
| CD29 | Protein | 0.14 | <1e-10 |
| LTB | Gene | -0.47 | <1e-10 |
| AQP3 | Gene | -0.26 | <1e-10 |
| IL7R | Gene | -0.25 | <1e-10 |
| KLRB1 | Gene | -0.23 | <1e-10 |
| CD27 | Gene | -0.21 | <1e-10 |
| MAL | Gene | -0.20 | <1e-10 |
| GZMH | Gene | 0.58 | <1e-10 |
| NKG7 | Gene | 0.55 | <1e-10 |
| FGFBP2 | Gene | 0.53 | <1e-10 |
| GNLY | Gene | 0.49 | <1e-10 |
| ZEB2 | Gene | 0.43 | <1e-10 |
| CCL4 | Gene | 0.39 | <1e-10 |

**Supplementary Table 7: Genes and proteins associated with CNA populations for TB progression**. Correlations between intensity (resp. expression level) of selected proteins (resp. genes) and the per-cell neighborhood coefficient computed by CNA for TB progressor status. Simple linear models between expression levels and neighborhood correlation values per-cell were used to determine naive p-values for each of these associations.

| Attribute tested | Covariates |
|---|---|
| Age | Weight, # scars, Winter blood draw<br>TB progression |
| Height | Weight, % European ancestry, Male sex<br>Edu. below high school, Winter blood draw, TB progression |
| Weight | Age, Height, % European ancestry<br>Edu. below high school, TB progression |
| % European ancestry | Height, Weight, Edu. below high school<br>Winter blood draw, TB progression |
| # scars | Age, BCG scar, Male sex<br>TB progression |
| BCG scar | # scars, Male sex, Winter blood draw |
| BCG vaccine | - |
| Male sex | Height, # scars, BCG scar<br>TB progression |
| Edu. below high school | Height, Weight, % European ancestry |
| High SES | - |
| Low SES | - |
| Medium SES | - |
| Smoking status | - |
| Alcohol use | - |
| Underweight | - |
| Spring blood draw | Winter blood draw, TB progression |
| Winter blood draw | Age, Height, % European ancestry<br>BCG scar, Spring blood draw |

**Supplementary Table 8: Covariate control for TB phenotypes tested**. For each sample attribute analyzed using CNA and MASC, we list the sample attributes included as covariates in the analysis.

| Phenotype | CNA P | Var. Exp. | Clusters+ | Clusters- | CNA+ | CNA- | % Replic. | % Novel |
|---|---|---|---|---|---|---|---|---|
| Age | < 1.0e-06 | 47% | 67882 | 41228 | 51032 | 118976 | 68 | 44 |
| Winter | < 1.0e-06 | 26% | 0 | 90946 | 30826 | 49112 | 44 | 50 |
| Sex | 2.0e-06 | 28% | 17085 | 77483 | 28507 | 147172 | 65 | 35 |
| Ancestry | 3.9e-04 | 16% | NA | NA | 6114 | 37230 | NA | 100 |

**Supplementary Table 9: Survey of associations in TB dataset**. Four per-sample outcome variables from the TB dataset were found to have global associations using CNA, of which one did not have a global association by cluster-based association testing using MASC. Global P values from CNA were subjected to Bonferroni correction for the number of sample attributes in the survey. Global associations for each tested outcome by CNA are tabulated, along with the percent of outcome variance explained by CNA. The total number of cells in all expanded clusters identified by MASC is also shown, where each cluster is considered significant only after Bonferroni correction for the number of clusters in the analysis. Likewise, the total cells in all depleted clusters is shown. The number of cells whose neighborhoods were found to be positively correlated in abundance with the outcome by CNA at an FDR 5% threshold are shown, as well as the number of cells whose neighborhoods were found to be negatively correlated with the outcome by CNA at an FDR 5% threshold. Finally, to illustrate the scope of recapitulated and novel associations, for the phenotypes for which local association were found by both methods we show the fraction of all cells found to belong to expanded or depleted clusters that were also assigned to associated populations by CNA ("% Replicated") and the fraction of all cells assigned to associated populations by CNA that did not belong to expanded or depleted clusters ("% Novel").

| Name | Type | Correlation | P.value |
|---|---|---|---|
| CD8a | Protein | 0.25 | <1e-10 |
| CD4.1 | Protein | -0.09 | <1e-10 |
| CD27.1 | Protein | -0.35 | <1e-10 |
| CD28.1 | Protein | -0.40 | <1e-10 |
| GZMH | Gene | 0.55 | <1e-10 |
| NKG7 | Gene | 0.51 | <1e-10 |
| FGFBP2 | Gene | 0.48 | <1e-10 |
| GNLY | Gene | 0.46 | <1e-10 |
| CST7 | Gene | 0.39 | <1e-10 |
| GZMA | Gene | 0.39 | <1e-10 |
| CCL5 | Gene | 0.37 | <1e-10 |
| CCL4 | Gene | 0.34 | <1e-10 |
| KLRB1 | Gene | -0.43 | <1e-10 |
| LTB | Gene | -0.42 | <1e-10 |
| LDHB | Gene | -0.19 | <1e-10 |
| TCF7 | Gene | -0.19 | <1e-10 |
| CCR6 | Gene | -0.17 | <1e-10 |
| CD27 | Gene | -0.15 | <1e-10 |

**Supplementary Table 10: Genes and proteins associated with CNA populations for age**. Correlations between intensity (resp. expression level) of selected proteins (resp. genes) and the per-cell neighborhood coefficient computed by CNA for age. Simple linear models between expression levels and neighborhood correlation values per-cell were used to determine naive p-values for each of these associations.

| Name | Type | Correlation | P.value |
|---|---|---|---|
| CD183/CXCR3 | Protein | -0.11 | <1e-10 |
| CD194/CCR4 | Protein | 0.28 | <1e-10 |
| GATA3 | Gene | 0.28 | <1e-10 |
| KRT1 | Gene | 0.26 | <1e-10 |
| ANXA1 | Gene | 0.26 | <1e-10 |
| IL7R | Gene | 0.25 | <1e-10 |
| STAT1 | Gene | -0.04 | <1e-10 |
| IFNG | Gene | -0.08 | <1e-10 |
| LIMS1 | Gene | -0.27 | <1e-10 |
| TIGIT | Gene | -0.27 | <1e-10 |
| CD27 | Gene | -0.24 | <1e-10 |
| TBC1D4 | Gene | -0.19 | <1e-10 |
| CTLA4 | Gene | -0.17 | <1e-10 |

**Supplementary Table 11: Genes and proteins associated with CNA populations for winter blood draw**. Correlations between intensity (resp. expression level) of selected proteins (resp. genes) and the per-cell neighborhood coefficient computed by CNA for winter blood draw. Simple linear models between expression levels and neighborhood correlation values per-cell were used to determine naive p-values for each of these associations.

| Name | Type | Correlation | P.value |
|---|---|---|---|
| CD8 | Protein | 0.04 | <1e-10 |
| CD194/CCR4 | Protein | 0.47 | <1e-10 |
| CD62L | Protein | 0.24 | <1e-10 |
| CD244/2B4 | Protein | -0.52 | <1e-10 |
| CD29 | Protein | -0.21 | <1e-10 |
| CD4 | Protein | -0.11 | <1e-10 |
| LTB | Gene | 0.46 | <1e-10 |
| LDHB | Gene | 0.27 | <1e-10 |
| SELL | Gene | 0.25 | <1e-10 |
| NKG7 | Gene | -0.72 | <1e-10 |
| CCL5 | Gene | -0.70 | <1e-10 |
| GZMH | Gene | -0.70 | <1e-10 |
| GZMA | Gene | -0.70 | <1e-10 |
| GNLY | Gene | -0.61 | <1e-10 |

**Supplementary Table 12: Genes and proteins associated with CNA populations for European ancestry**. Correlations between intensity (resp. expression level) of selected proteins (resp. genes) and the per-cell neighborhood coefficient computed by CNA for European ancestry. Simple linear models between expression levels and neighborhood correlation values per-cell were used to determine naive p-values for each of these associations.

#### Supplementary Figures

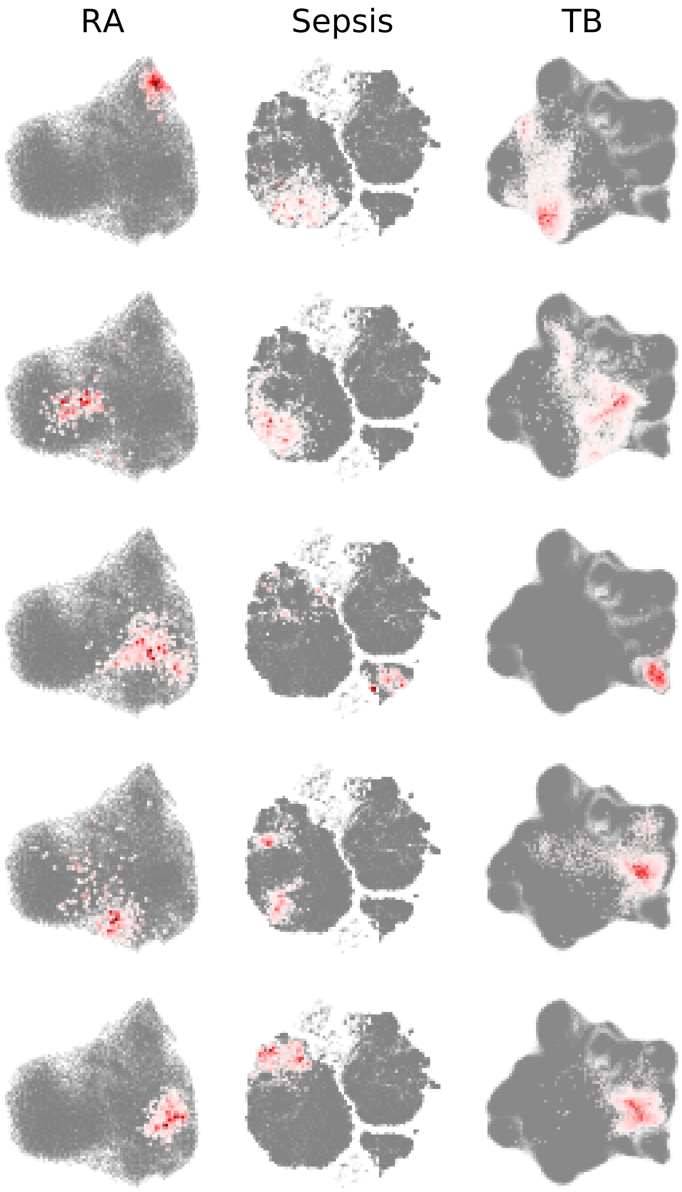


**Supplementary Figure 1: Example neighborhoods in three real datasets**. Each column shows five example neighborhoods from the indicated dataset among the three datasets analyzed in the paper. Cells are colored according to their degree of belonging to the neighborhood. In each case, only the "bulk" of each neighborhood is shown: that is, only cells with belonging above a certain threshold are shown, with the threshold set such that 70% of the mass of the neighborhood is included.

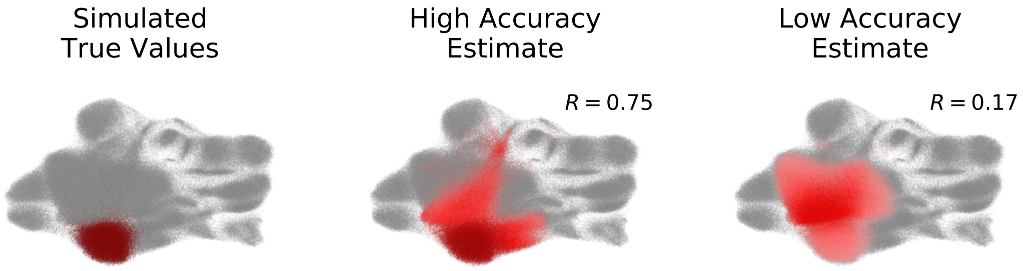


**Supplementary Figure 2: Accuracy, illustrated**. Consider a simulated signal of the cluster abundance type, where true values per cell are assigned to be 1 for all cells in the selected cluster and 0 elsewhere (**Left**). Our resulting per-sample values are the fraction of that sample's cells from the selected cluster. A model estimate of this causal population with high accuracy closely approximates the true direction and degree of abundance association to the simulated per-sample values across transcriptional space (**Center**). A model estimate with lower accuracy does not accurately identify the true causal population (**Right**).

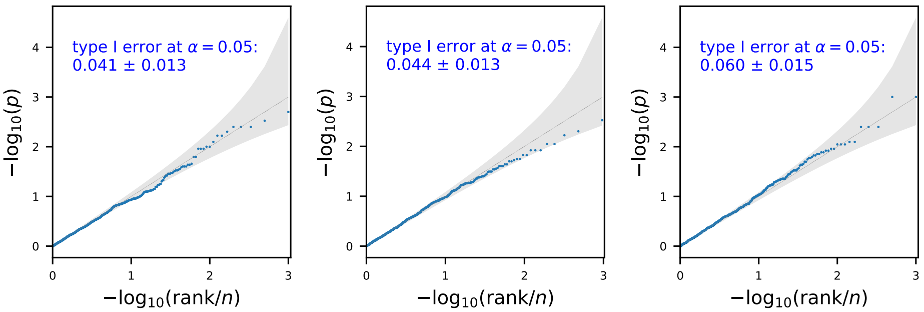


**Supplementary Figure 3: Calibration of CNA**. P values from 1,000 trials when CNA is conducted with simulated per-sample outcomes of: patient age values permuted randomly across the dataset (**Left**), or patient age values permuted within batch, to test calibration under moderate batch effects (**Middle**). We then tested calibration under extreme batch effects (**Right**) simulating outcomes with a value of 1 for all samples in a given batch and 0 otherwise, across 1,000 randomly chosen batches.

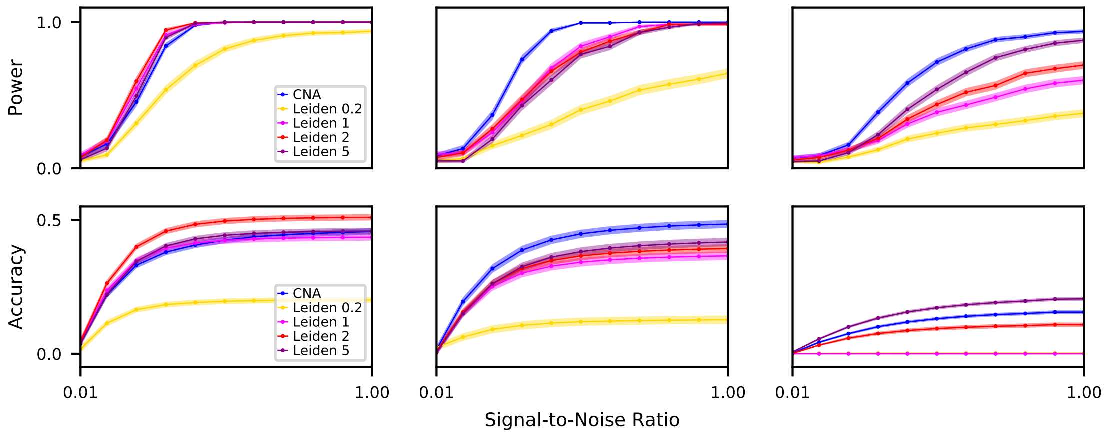


**Supplementary Figure 4: Power and accuracy shown separately by clustering resolution**. Three signal types – causal clusters, global gene expression programs, and cluster-specific gene expression programs – were simulated, corresponding to the left, middle and right columns, respectively. For each signal type, we show (**top**) the relative power of CNA versus a cluster-based approach across a range of signal:noise ratios, and (**bottom**) the relative accuracy of CNA versus a cluster-based approach across a range of noise levels.

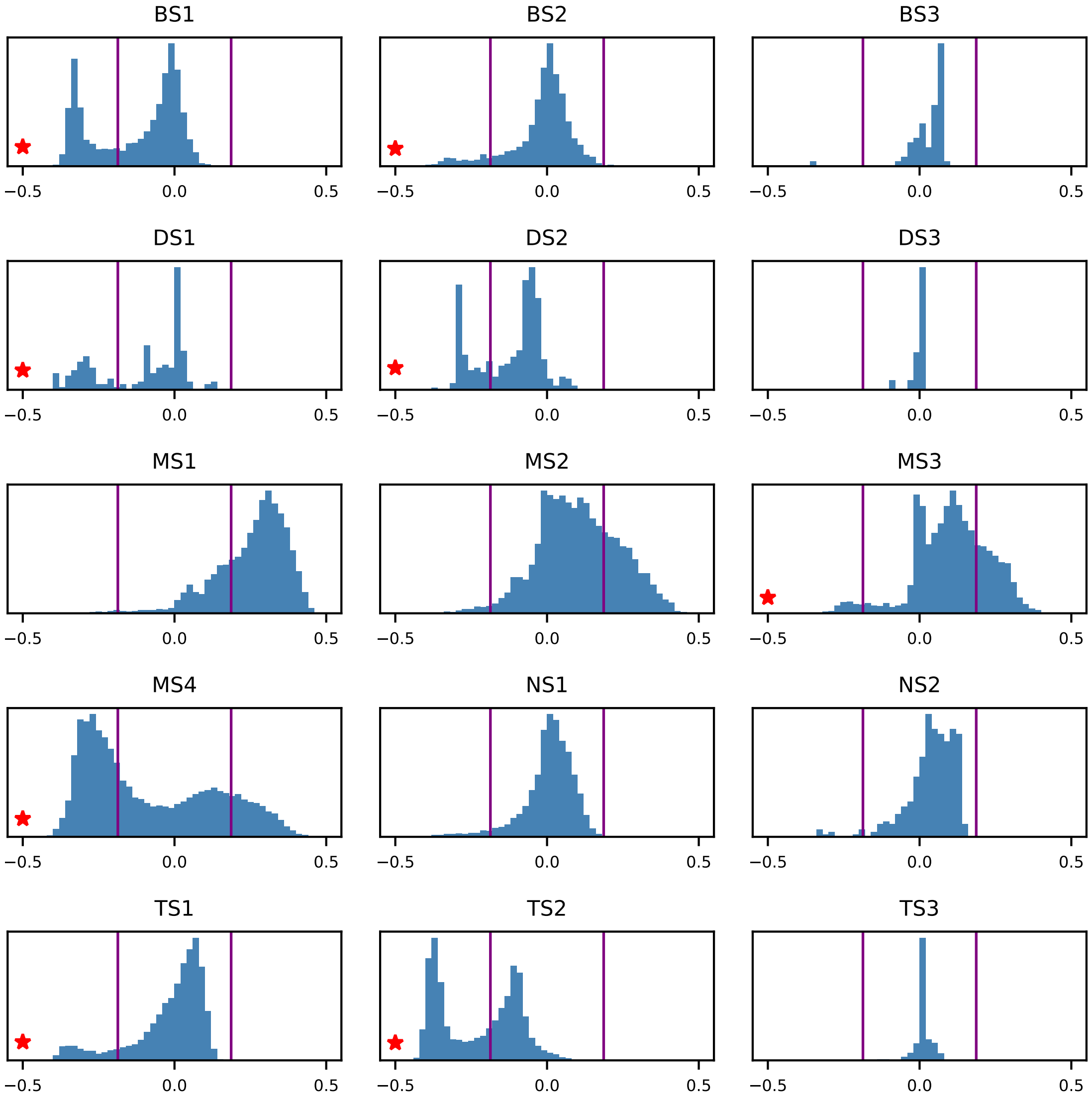


**Supplementary Figure 5: Within-cluster heterogeneity in sepsis dataset**. Histograms of neighborhood coefficients for the sepsis versus no-sepsis phenotype, within each published cluster for this dataset. The abundance correlation thresholds beyond which the anchor cell for each neighborhood was assigned to an expanded-in-sepsis population (right) or a depleted-in-sepsis population (left) are marked with purple vertical lines. Clusters containing sub-populations with distinct abundance associations to the sepsis phenotype are starred.

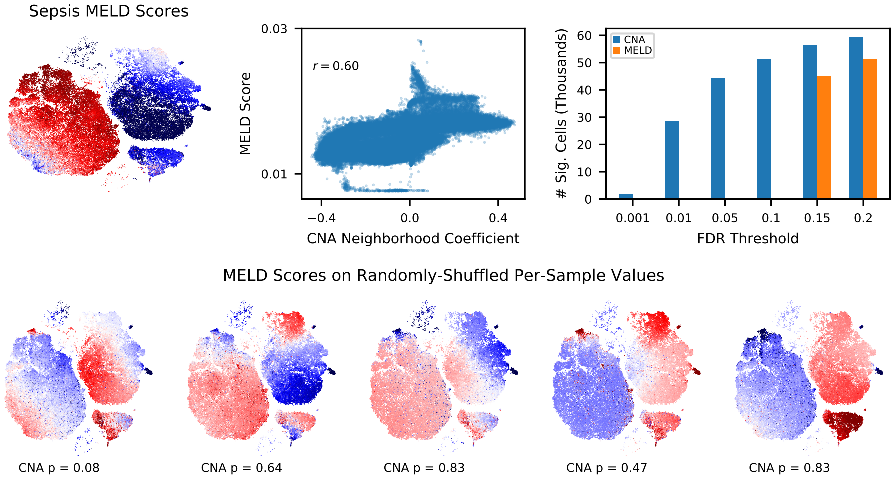


**Supplementary Figure 6: Sepsis dataset analysis with MELD**. We first applied the MELD algorithm to the sepsis dataset using the true sepsis vs non-sepsis sample attribute values. Per-cell scores from MELD are shown in tSNE space (**Top Left**). MELD scores were correlated with the neighborhood coefficients from CNA (Pearson R=0.60, **Top Middle**). MELD does not report a global significance metric, so we sought to determine whether the MELD score pattern from the sepsis attribute was more striking than MELD score patterns from null comparator attributes. We randomly permuted the sample sepsis vs non-sepsis attribute values five times and examined the resulting MELD scores, which appear to have nontrivial structure across transcriptional space (**Bottom Row**). For reference, we also ran CNA on these null attributes and all CNA global p-values were sub-significant. To evaluate the local significance of MELD scores per-cell from the true sepsis attribute, we applied a permutation-based approach identical to the one used by CNA to assess significance per-neighborhood. None of the individual per-cell MELD scores were significant at FDR¡0.05 (**Top Right**). More specifically, we generated 500 null distributions of MELD scores each resulting from a random permutation of the sample sepsis case-control labels. We then used these null instantiations to estimate the false discovery rate for MELD scores. We plot the number of cells (for MELD) and neighborhoods (for CNA) that pass an increasing FDR threshold.

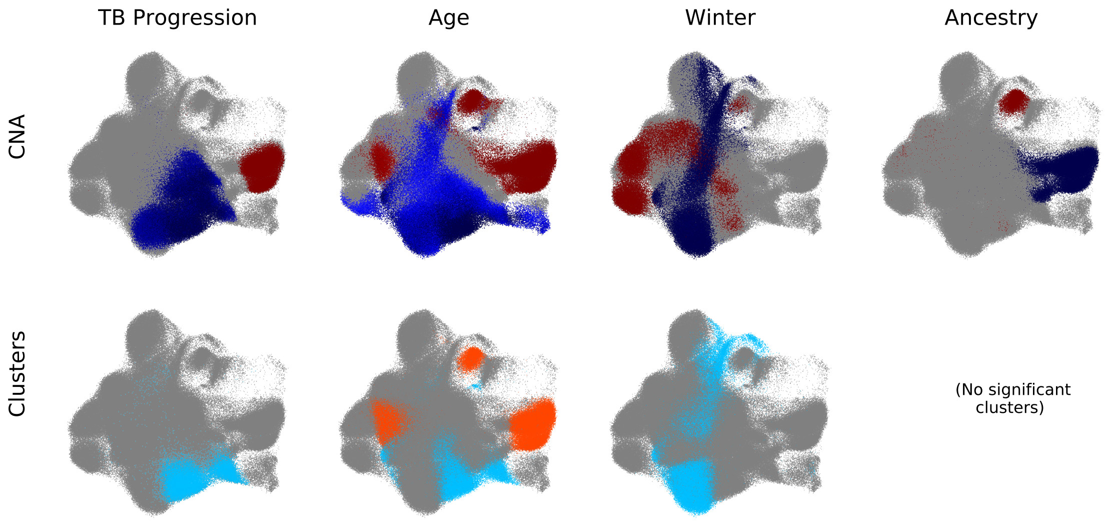


**Supplementary Figure 7: Contrasting cell populations implicated by CNA compared to cluster-based analysis**. (**Top**) Cell populations implicated by CNA as depleted (blue) or expanded (red) in association with each sample attribute, arrayed left to right. (**Bottom**) Cell populations implicated by cluster-based analysis as depleted (blue) or expanded (orange) in association with each sample attribute. We note that although our cluster-based analysis did not recover significant local associations for ancestry, the cluster-based analysis in the original study did; this difference is primarily due to the two analyses using different ways of choosing which covariates to include as well as the more aggressive multiple-testing correction across phenotypes employed in our analysis.
